# Predicting functional topography of the human visual cortex from cortical anatomy at scale

**DOI:** 10.1101/2025.11.27.690210

**Authors:** Fernanda L. Ribeiro, Robert Satzger, Felix Hoffstaedter, Christian Bürger, Peer Herholz, David Linhardt, Noah C. Benson, D. Samuel Schwarzkopf, Alexander M. Puckett, Steffen Bollmann, Martin N. Hebart

## Abstract

Topographic organization, whereby neighboring cortical locations encode neighboring features in sensory or cognitive space, is a fundamental principle of brain function. Existing approaches for obtaining individual-specific topographic maps either require resource-intensive functional neuroimaging or, when relying on population atlases, lack precision for individual-level inference. Here, we introduce *deepRetinotopy toolbox*, a deep learning-based application for predicting the functional topographic organization of human visual cortex from cortical anatomy alone. *DeepRetinotopy toolbox* produces accurate retinotopic maps across diverse experimental conditions, imaging sites, and scanner types. We demonstrate how predicted maps can be utilized to automatically generate individual-specific visual area boundaries, overcoming common biases in manual annotations. Finally, we applied our method to 11,060 anatomical scans, which allowed us to quantify age-related changes in the functional organization of visual cortex predictable from anatomy alone, underscoring the method’s broad utility for scalable, anatomy-based functional brain mapping.

## Introduction

Topographic organization is a fundamental principle of information processing in the human brain^1^. A topographically organized brain region represents information in a spatially orderly manner, with adjacent locations in the brain responding to adjacent features along some physical or cognitive dimension. For example, in the visual cortex, neurons encode spatial location in the visual field in a *retinotopic* format, while in the somatosensory cortex, neurons encode spatial location along the body skin in a *somatotopic* format. This spatially cohesive representation of sensory information extends beyond the visual cortex and other primary sensory areas^2,3^ and may even serve as the structural scaffold for higher-order cognitive processes^4,5^. Therefore, understanding these topographic maps is essential for advancing basic neuroscience and investigating brain-behavior relationships.

Despite their fundamental importance, current methods for understanding topographic organization in humans face important limitations that hinder large-scale studies and clinical applications. For example, acquiring high-quality functional MRI (fMRI) data to estimate topographic organization in the visual cortex is time-consuming and costly, and requires considerable expertise in stimulus design and specialized data analysis. Alternatively, researchers often rely on atlases of areal boundaries^6–8^ and population-average topographic maps^9^. Still, these do not capture the substantial individual differences^9,10^ that are crucial for understanding brain-behavior relationships^11–13^. This creates a fundamental dilemma: empirical mapping is typically too resource-intensive for large-scale applications, while atlas-based approaches sacrifice much of the individual specificity essential for strong individual-level inference of behavior.

The tight coupling between topographic organization and cortical anatomy in the visual cortex offers a potential solution to this dilemma. Previous work has demonstrated that cortical folding is a useful predictor of topographic organization in early visual cortex^8,15–17^. Proof-of-concept work (deepRetinotopy-beta)^18^ has demonstrated that geometric deep learning models can not only learn the complex structure-function relationship of the visual cortex but also capture fine-grained individual differences in retinotopic organization^10^. However, this approach has remained limited by requiring specific kinds of anatomical data that are not commonly available, ultimately preventing a comprehensive generalizability assessment and application to the vast number of existing anatomical datasets. In addition, accessible tools for similar approaches have been lacking, hindering widespread adoption of similar techniques.

To address these challenges, here we introduce *deepRetinotopy toolbox*, a robust and accessible framework that realizes this potential by enabling large-scale topographic mapping of the visual cortex by leveraging common anatomical MRI scans. We benchmark and comprehensively assess the generalizability of our approach using multiple publicly available fMRI datasets acquired under diverse experimental conditions, including varying imaging sites, scanner types, and visual stimuli protocols used for retinotopic mapping. Our findings demonstrate that *deepRetinotopy toolbox* can accurately predict retinotopic organization in new datasets from cortical anatomy alone. Moreover, we demonstrate that these predicted maps can replace empirical measurements to automatically segment and differentiate early visual areas. Finally, we demonstrate the potential of this approach by retrospectively applying it to thousands of brain scans, revealing that age-related differences in primary visual cortex organization^19^ can be predicted from cortical anatomy. This finding demonstrates the new potential of historical datasets with hundreds of thousands of existing anatomical scans to obtain new insights into structure-function coupling in the visual cortex, how it may change during development, or how it may vary between populations. Beyond retinotopy, this work establishes a general framework for leveraging structure-function relationships to predict topographic organization in other parts of the brain. Our toolbox requires minimal data and provides a user-friendly command line interface, enabling scalable deployment across diverse computing environments for broad adoption by the neuroscience community.

## Results

### Toolbox overview

To obtain representations of the visual field in the human brain, that is, retinotopic maps, researchers typically rely on the acquisition of functional MRI data from participants while they look at visual stimuli that are temporally and spatially controlled. These visual stimuli commonly consist of rotating wedges, expanding and contracting rings, and sweeping bars. The analysis of the fMRI signals elicited by such stimuli enables the estimation of the visual field location that is preferentially encoded by each responsive voxel^20^. However, because these retinotopic maps are highly correlated with the underlying cortical anatomy^17,21–24^, the folds and curves of individual brains contain rich information to predict where different parts of the visual field are represented. Specifically, cortical curvature patterns serve as anatomical landmarks exhibiting a high degree of consistency across individuals, enabling computational models to capture shared topographic organization from anatomy alone^25^. Our *deepRetinotopy toolbox* introduces a general, dataset-agnostic framework that exploits this structure-function relationship to predict retinotopic organization from cortical anatomy.

Our toolbox integrates standard neuroimaging software for anatomical MRI data preprocessing and pre-trained deep learning models for predicting topographic maps of the visual cortex at the level of individuals (Figure 1a). These topographic maps include visual field maps (polar angle and eccentricity) which capture the spatial organization of visual field representations. In addition, the population receptive field (pRF) size map quantifies the extent of the visual field that elicits a response in a given voxel, which can be estimated as an additional parameter from fMRI signals in response to visual stimuli^20^. Throughout this paper, we use ’retinotopic maps’ to refer to all map types. Our deep learning models were trained using a subset (n = 161) of the Human Connectome Project (HCP) 7T Retinotopy dataset^26^ (see *Methods* section *Model training and selection*).

**Figure 1.**
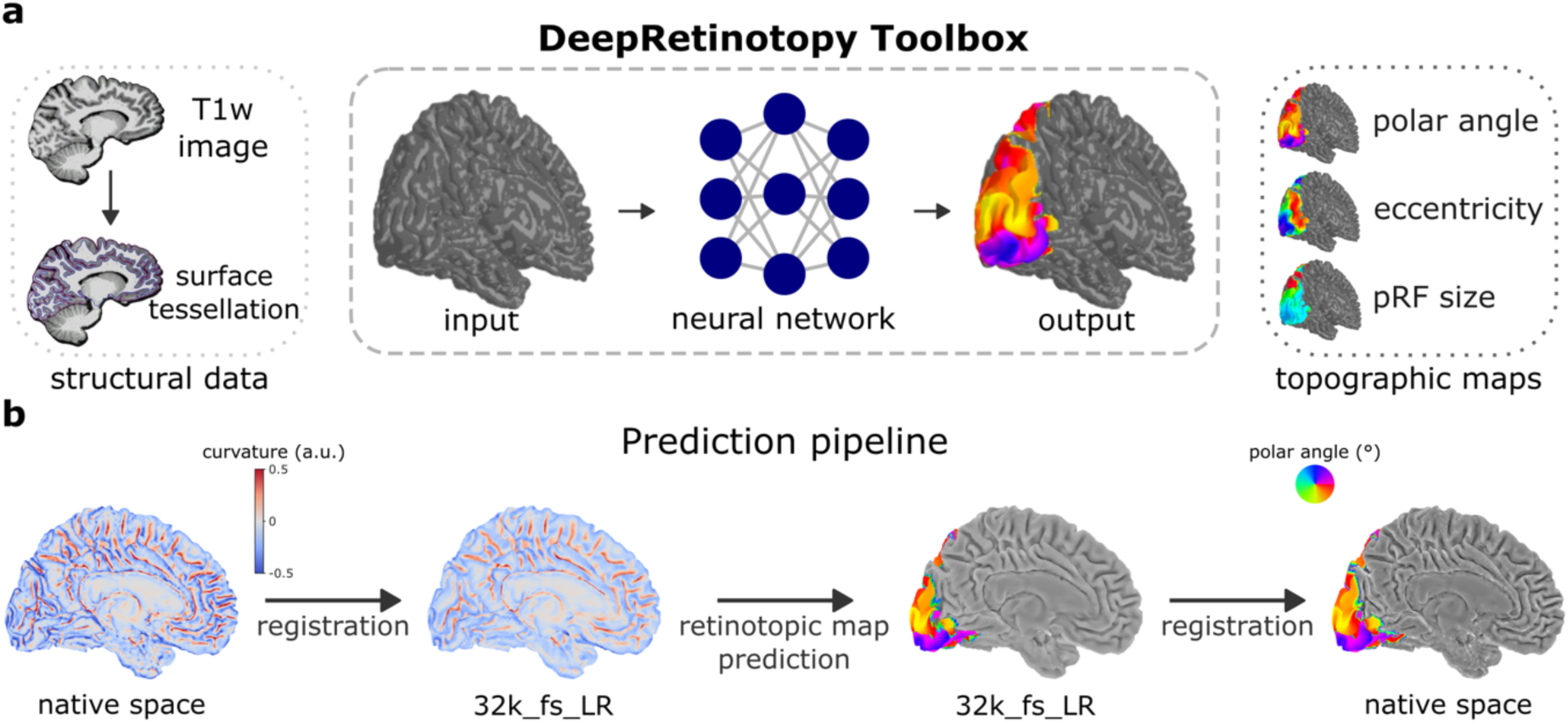
DeepRetinotopy toolbox: an application for predicting retinotopic organization from brain structure. **a,** DeepRetinotopy toolbox integrates standard neuroimaging software for anatomical MRI data preprocessing (left) and pre-trained deep learning models (middle) for predicting retinotopic maps at the individual level (right). **b,** The prediction pipeline requires FreeSurfer-reconstructed cortical meshes derived from a T1w structural MRI image as input. Note that our toolbox can perform both the preprocessing and prediction steps, or users can provide preprocessed data directly. The prediction pipeline consists of registering curvature maps from the native space to a template space (32k_fs_LR). These registered curvature maps are then used as input for retinotopic map prediction using pre-trained models. Finally, predicted maps are then registered to the individual’s native space.

The toolbox is modular and consists of three main processing steps (Figure 1b). First, it processes the input data, that is, cortical meshes reconstructed from structural MRI images (T1w) using *FreeSurfer*, to estimate mean curvature maps and bring them to a common surface space. Next, the pre-trained models predict individual-specific retinotopic maps from input data. Finally, predicted maps are resampled to the native space from the anatomical data. These modules are combined in an easy-to-use command line interface that is compatible with common neuroimaging pipelines^27,28^ and the brain imaging data structure (BIDS)^29^. The toolbox is packaged into software containers^30,31^ to enable reproducible and scalable processing and can be easily used on Neurodesk^31^ as a module or in any computational environment with a compatible container runtime.

### Benchmarking: predicting retinotopic maps of the human visual cortex from underlying anatomy

To benchmark our method, we compared our approach with two other alternative methods for predicting retinotopic maps of the visual cortex from anatomy. These include: (1) a model with a similar architecture as ours that requires both curvature and myelin maps as input features^18^, which we refer to as **deepRetinotopy-beta**, and (2) an anatomical template of retinotopy^15^, which we refer to as **Benson2014**. DeepRetinotopy-beta demonstrated very good performance on the HCP test set using both curvature and myelin maps. However, it has not been applied to different datasets, mainly due to specific data requirements. For example, myelin maps are derived from a combination of MRI image contrasts^32^ that are not collected by default in most studies, which prevents the method from being applied to most existing and future datasets. Benson2014 consists of an anatomical template of retinotopy to which individual participant structural data is registered, yielding retinotopic map predictions at the level of the individual^15^. For qualitative assessment, Figure 2a shows retinotopic maps in early visual areas (V1-3) from a single participant in the HCP test set. Both *deepRetinotopy* and deepRetinotopy-beta were able to generate more realistic polar angle and pRF size maps than Benson2014, while eccentricity maps were qualitatively similar across methods. Figure 2b shows the average correlation between predicted and empirically derived maps across all participants in the HCP test set, with the gray shaded area representing the noise ceiling (see *Methods* section *Benchmarking*). *DeepRetinotopy* performance was on par with deepRetinotopy-beta across retinotopic maps, even though it only relied on the individual’s curvature map as an input feature. Moreover, Benson2014 is considerably worse for predicting eccentricity and pRF size. Supplementary Figure 1 shows consistent findings focusing on error estimation. Taken together, this shows that *deepRetinotopy* can predict retinotopic organization at the level of individuals, only based on T1w MRI images.

**Figure 2.**
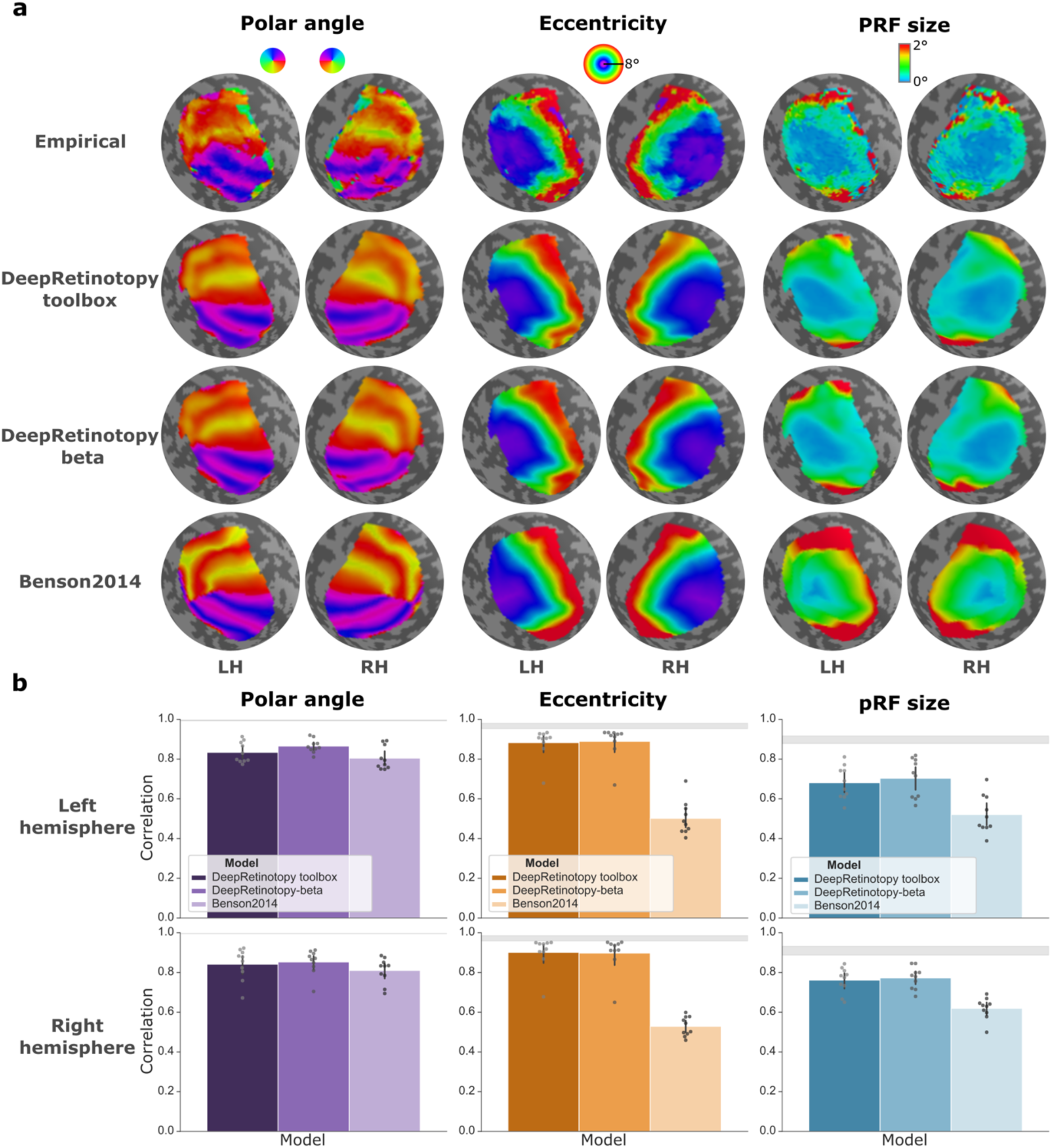
Benchmarking models for predicting retinotopic maps of the visual cortex from underlying anatomy. **a**, Polar angle, eccentricity, and pRF size maps from left (LH) and right (RH) hemispheres are shown from a representative participant in the HCP test set (#680957). **b**, Bar plots represent the average correlation between predicted and empirically derived maps across all participants in the HCP test set (n = 10). Correlation score was determined as either the Pearson correlation (for eccentricity and pRF size maps) or the circular correlation (polar angle maps). Retinotopic maps were vectorized, and only vertices within early visual areas (V1-3) and above a 10% variance explained threshold were used to estimate the correlation. Error bars correspond to the 95% confidence interval. The gray shaded area represents the noise ceiling, i.e., the 95% confidence interval of the square root of the Spearman-Brown corrected variance explained between split-half pRF fits.

### Generalizability: prediction performance across diverse datasets

To demonstrate the general utility of *deepRetinotopy* and allow retinotopic mapping from brain anatomy for the tens-of-thousands of individuals for whom anatomical scans are available, our method must generalize to new datasets. Here, we broadly assessed the generalizability of our approach using datasets acquired across different experimental conditions. These new datasets vary in several ways, including imaging sites, scanner types, data resolution, visual stimuli, and encoding models used for reconstructing retinotopic maps (Figure 3a). Cross-dataset comparisons^33^ (see also Supplementary Figure 2-4) indicate that empirically estimated retinotopic maps—especially eccentricity and pRF size—vary systematically with experimental factors such as fMRI preprocessing and visual stimuli. These variations, however, occur in predictable ways. For example, pRF size tends to be overestimated in foveal regions with sweeping bar stimuli compared to a combination of visual stimuli^34^ (i.e., wedge, ring, and bars). These parameter estimation differences are, therefore, an intrinsic consequence of how retinotopic mapping is commonly performed. Thus, we expected deviations in predicted *versus* empirically derived maps to occur in systematic ways. Where such deviations occur in predictable ways, they reflect differences in the experimental design used in the training data rather than limitations of our approach.

**Figure 3.**
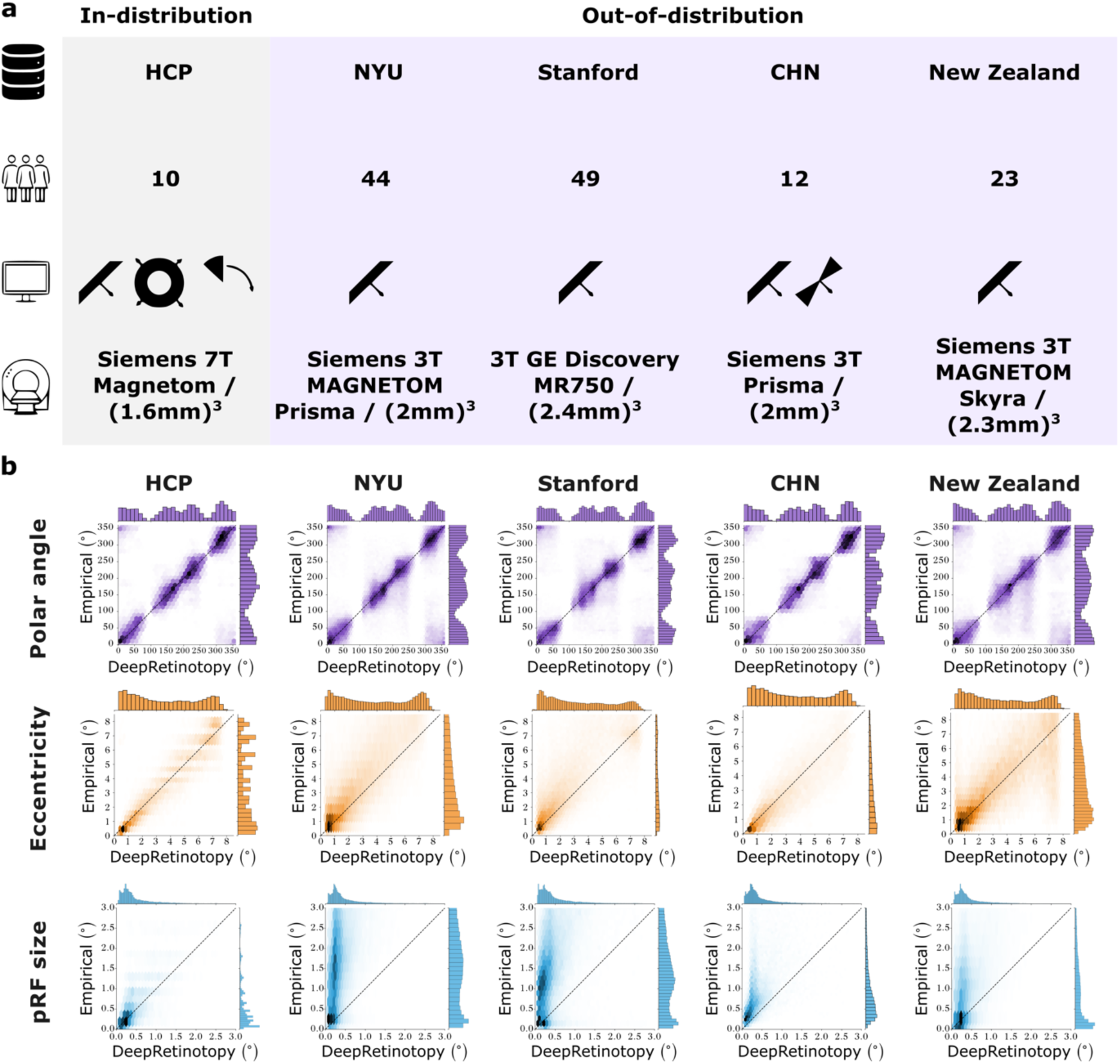
Cross-dataset generalizability. **a**, Diagram of the main characteristics of the datasets used to assess model generalizability. From top to bottom, we show: dataset name/acronym, sample size, visual stimuli used for retinotopic mapping experiments, scanner type, and fMRI data resolution. In grey, we highlight the HCP test set and, in purple, four distinct new datasets. **b**, Empirically derived and predicted maps were vectorized, filtered to include only vertices from V1-3 and above a variance explained threshold of 10% of the empirical data, concatenated across participants, and represented as hexbin plots. In each plot, empirically derived parameters are shown along the y-axis and the predicted ones along the x-axis. The top row shows polar angle values, while the middle and bottom rows show the equivalent plots for eccentricity and pRF size maps. Black diagonal lines illustrate the ‘perfect’ match between empirically derived and predicted parameters.

Figure 3b shows the empirically estimated *versus* predicted parameters across vertices within early visual areas and participants from each dataset. We found that predicted polar angle maps (Figure 3b, top) were highly accurate across datasets [HCP: r_LH_ = 0.834; NYU: r_LH_ = 0.691; Stanford: r_LH_ = 0.674; CHN: r_LH_ = 0.810; New Zealand: r_LH_ = 0.637; statistical significance was tested using a spatial autocorrelation-preserving null model (‘spin test’)^35^ and all results were statistically significant (p<.001)], indicating the generalizability of our approach for capturing this topographic organization. Similarly, eccentricity predictions (Figure 3b, middle) showed high correlations across sites (HCP: r_LH_ = 0.882; NYU: r_LH_ = 0.759; Stanford: r_LH_ = 0.690; CHN: r_LH_ = 0.864; New Zealand: r_LH_ = 0.728). In contrast, while prediction performance for pRF size maps (Figure 3b, bottom) remained high for the HCP test set (r_LH_ = 0.681), as expected, performance declined across new datasets, consistent with known cross-dataset variability in this parameter^33^, with correlations ranging from 0.485 (CHN) to 0.687 (NYU). Specifically, *deepRetinotopy* predicted smaller pRF sizes relative to the other datasets due to choices in the experimental design of the training data (i.e., the HCP dataset; Supplementary Figure 4), including visual stimuli. Supplementary Table 1 shows the complete set of results for both hemispheres. Together, this demonstrates the high generalizability of retinotopic mapping predictions using *deepRetinotopy toolbox* to other datasets.

### Automated visual area segmentation

Having demonstrated that *deepRetinotopy* generalizes across datasets, we next aimed to highlight potential applications of the approach. Accurate delineation of individual-specific visual areas, such as V1, V2, or V3, is necessary for most studies of the visual cortex that aim to localize functional responses, investigate clinical deficits precisely, and determine biological correlates of individual differences in perception^12,36^. Traditionally, this first requires the acquisition of retinotopic mapping data, followed by laborious manual annotation by expert annotators. However, this approach is not only time-consuming but also introduces idiosyncrasies related to individual annotators or scanning sites. Here, we demonstrate how *deepRetinotopy* enables automated visual area segmentation from standard anatomical data alone, thereby overcoming the long-standing reliance on manual delineation and opening new possibilities for large-scale population studies. We propose a fully automated approach by combining our toolbox with the Bayesian model of retinotopy^9^ to produce individual-level visual area boundaries (Figure 4a).

**Figure 4.**
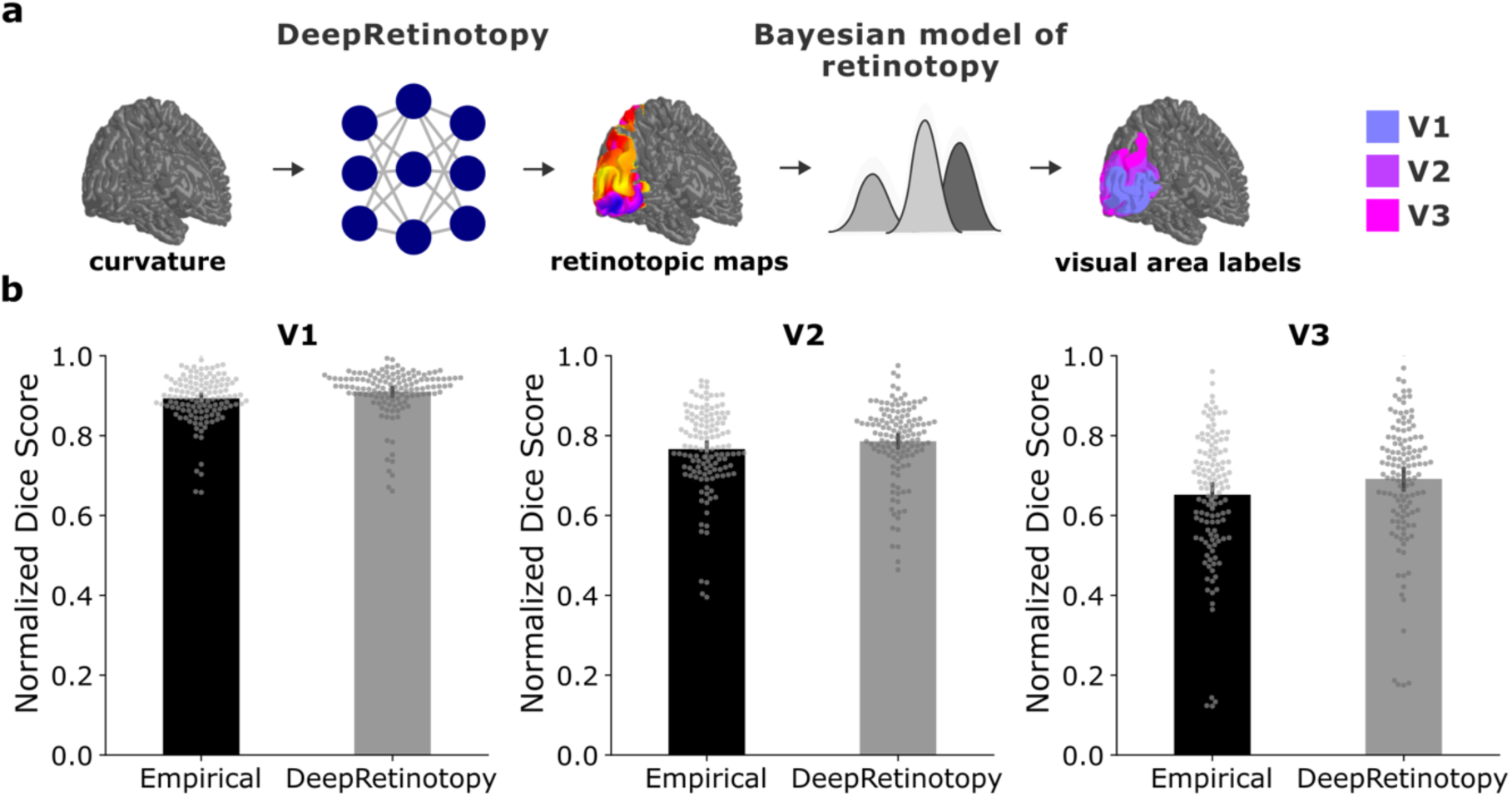
Automated visual area segmentation. **a**, Diagram shows our automated visual area segmentation pipeline, in which *deepRetinotopy*’s predicted retinotopic maps are used in combination with the Bayesian model of retinotopy to infer early visual area boundaries. **b**, Segmentation performance is shown across visual areas. Performance was estimated as the degree of overlap (Dice) between manually drawn and automatically generated early visual area labels, for which data from both hemispheres were combined. Then, each individual’s Dice scores were normalized by the corresponding mean Dice score between all pairs of manual annotations derived from four expert annotators. Error bars correspond to the 95% confidence interval.

The utility of this approach depends critically on two aspects: (1) whether predicted maps can substitute for empirical measurements for automated visual segmentation; and (2) whether automatically derived segmentation can substitute manual work (the gold standard). Therefore, we validated our approach by comparing automatically derived labels using either the empirically estimated or the predicted retinotopic maps with manually drawn labels by four expert annotators (Figure 4b and Supplementary Figure 5). Comparing automated segmentation derived from *deepRetinotopy* predicted maps with segmentation from empirically measured maps, we found slightly better segmentation performance for predicted maps (***V1***: mean normalized Dice score_DeepRetinotopy_ = 0.91 ± 0.06, mean normalized Dice score_empirical_ = 0.89 ± 0.06, p < .01; ***V2***: mean normalized Dice score_DeepRetinotopy_ = 0.79 ± 0.10, mean normalized Dice score_empirical_ = 0.77 ± 0.10, p < .01, ***V3***: mean normalized Dice score_DeepRetinotopy_ = 0.69 ± 0.16, mean normalized Dice score_empirical_ = 0.65 ± 0.16, p < .01; statistical significance was tested using a two-tailed paired t-test). For hemisphere-specific effects, see Supplementary Figure 6. Comparing these results to the gold standard of manually drawn labels, our results show that segmentation performance was excellent for V1 (normalized Dice score approaches 1), good to very good for V2, and acceptable to good for V3. Altogether, these results indicate that *deepRetinotopy* maps can replace empirical measurements to generate individual-specific early visual area boundaries, and that the automated segmentation can fully replace manual work for segmenting V1 but may need further improvement for reaching expert-level segmentation in V2 and V3.

### *DeepRetinotopy* enables large-scale investigation of individual differences in visual cortex organization

The demonstrated generalizability of *deepRetinotopy* across datasets makes it possible to apply the toolbox retrospectively to the tens of thousands of individuals with existing anatomical brain scans, thus enabling large-scale investigations of individual differences in retinotopic organization. To illustrate the scope of the approach, we ran our toolbox on 11,060 brain scans to uncover age-related variation in visual field representations. Beyond the well-established effects of cortical magnification—the cortical overrepresentation of central vision—polar angle asymmetries have been largely overlooked, until recently^38,39^. These asymmetries in cortical representation of the horizontal and vertical meridians vary between children and adults^19^ (Figure 5a) and are directly mirrored in behavior^11,40^. Here, we hypothesized that if age-related differences in cortical polar angle asymmetries emerge from changes in local cortical geometry, *deepRetinotopy* should detect them, thus providing an ideal test case for our approach.

**Figure 5.**
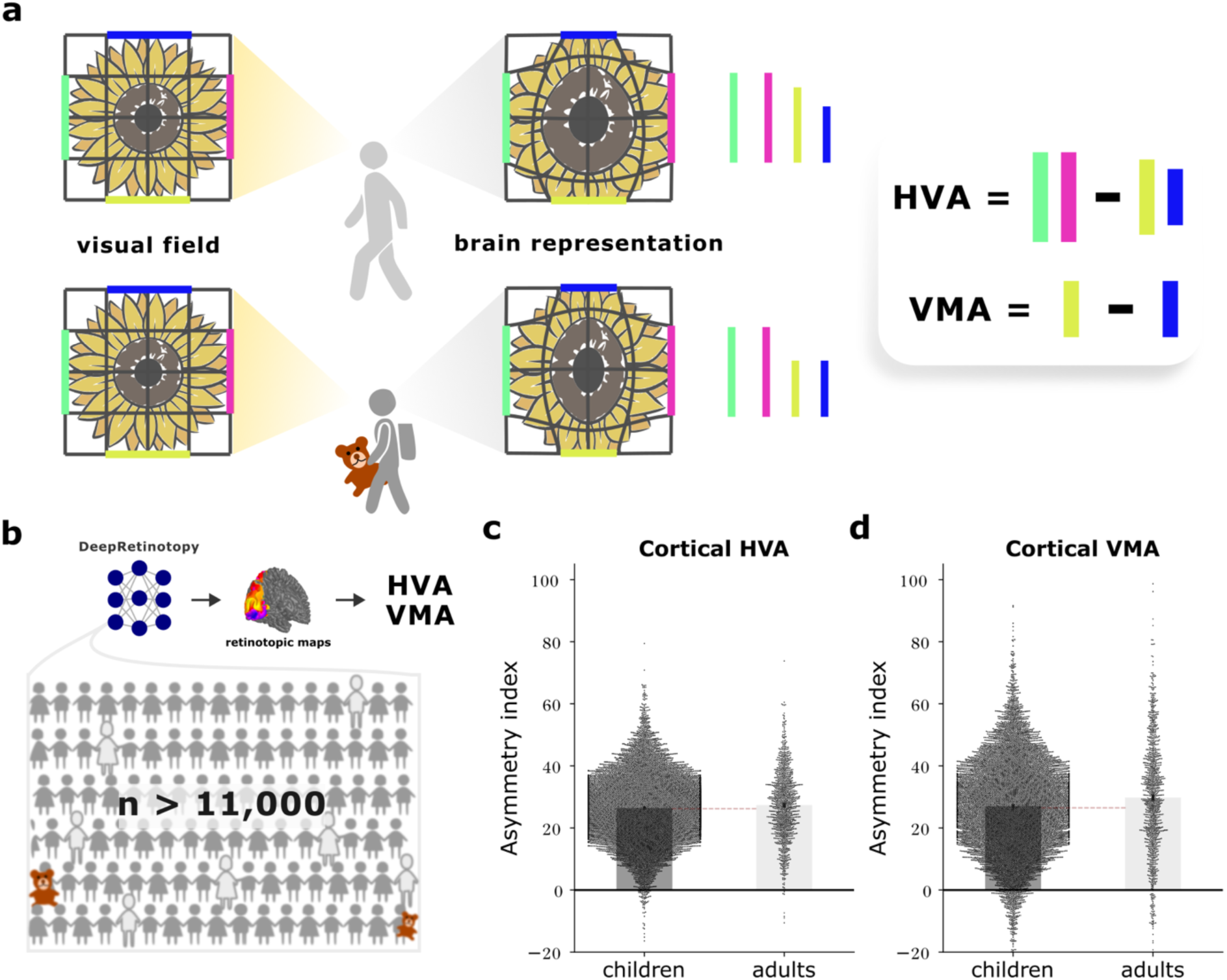
Cortical horizontal-vertical anisotropy (HVA) and vertical-meridian asymmetry (VMA) differences between children and adults. **a**, Diagram shows how the internal brain representation (middle) of the visual field (left) varies across eccentricity and polar angle. A greater surface area is devoted to central *versus* peripheral vision (note the expanded disc florets *versus* contracted ray florets in the brain representations). Moreover, more cortical surface area is dedicated to representing the horizontal meridian than the vertical meridian (HVA; represented as the difference between the combined pink and green edges, and yellow and blue edges), and more surface area is dedicated to representing the lower than the upper vertical meridian (VMA; represented as the difference between the yellow and blue edges). Cortical polar angle asymmetries also vary between adults (top) and children (bottom). Specifically, while HVA is similar between groups, children have reduced cortical VMA. **b**, We applied *deepRetinotopy* to over 11,000 brain scans and estimated both cortical VMA and HVA. Bar plots show the magnitudes of HVA (**c**) and VMA (**d**) indices for children and adults. Individual data points are also shown. Error bars correspond to ± standard error. Statistical comparisons using two-tailed independent samples t-test revealed significant group difference in cortical VMA (t(11,058) = 5.72, p < 0.001, d = 0.18) as well as cortical HVA (t(11,058) = 2.74, p < 0.01, d = 0.09).

To test this hypothesis, we first validated our predictions by replicating the previously described asymmetries in V1 surface area allocation of different portions of the visual field^39^. We found more cortical surface area dedicated to representing the horizontal meridian than the vertical meridian (horizontal-vertical anisotropy or HVA) and more surface area dedicated to representing the lower than the upper vertical meridian (vertical-meridian asymmetry or VMA) in both empirically derived and predicted retinotopic maps (Supplementary Figure 7). Moreover, these findings generalized to a large-scale dataset, the complete HCP Young Adult dataset^41^. These results support the notion that our approach can capture the asymmetries in visual field representation in V1.

To determine the potential of *deepRetinotopy* to uncover age-related variation in visual field representation, we applied our toolbox to a large-scale developmental dataset. Specifically, while cortical HVA is similar between adults and children, children have smaller cortical VMA^19^ (Figure 5a). To investigate whether cortical VMA emerges, at least in part, due to changes in local cortical geometry, we applied *deepRetinotopy* to the ABCD baseline cohort^42^ (n = 9,947; age 9-11) and compared it with the HCP Young Adult dataset (n = 1,113, age 22-36; Figure 5b). We operationalized these asymmetries by computing an asymmetry index given as the difference between the surface area of two wedge-ROIs (horizontal *versus* vertical meridian, and lower vertical *versus* upper vertical meridian) divided by the mean combined surface area (see *Methods* section *Estimating V1 surface area dedicated to sample different portions of the visual field* for further details). As in previous work, our approach was able to uncover a group difference in cortical VMA (t(11,058) = 5.72, p < 0.001, d = 0.18, two-tailed independent samples t-test; Figure 5d). Moving beyond previous findings, we found a small but significant difference in cortical HVA between groups (t(11,058) = 2.74, p < 0.01, d = 0.09, two-tailed independent samples t-test; Figure 5c). Importantly, these findings indicate that asymmetries in how the visual cortex samples different meridians can emerge from changes in local cortical geometry. More broadly, these findings showcase the application of *deepRetinotopy* to large-scale anatomical datasets for making novel discoveries about individual differences in functional topographic organization.

### Computational Efficiency

Lastly, we evaluated the computational efficiency of *deepRetinotopy* for large-scale studies. We benchmarked inference time and peak memory usage across two computing environments (Table 1). In both cases, retinotopic map prediction was performed on CPU only. Prediction of all three retinotopic maps for both hemispheres required approximately 250 seconds on a single-thread, high-performance computing cluster job with 64 CPU core (AMD EPYC 7742). The peak RAM usage was ∼1.4 GB. These modest computational requirements enable large-scale application to existing neuroimaging datasets without specialized hardware.

**Table 1.** Measures of computational efficiency in two computing environments.

| Computing environment | System 1 | System 2 |
| --- | --- | --- |
| Hardware | Apple M4 Max | AMD EPYC 7742 64-Core Processor |
| Operating System | MacOS Sequoia 15.6.1 | Debian GNU/Linux 11 (bullseye) |
| Container platform | Docker | Singularity/Apptainer |
| Inference time |  |  |
| All maps | 209.63s | 253.16s |
| Single map | 72.58s | 96.34s |
| Peak RAM |  |  |
| All maps | 1.5GB | 1.4GB |
| Single map | 1.5GB | 1.4GB |

## Discussion

A fundamental goal in neuroscience is to uncover how individual differences in brain organization relate to individual differences in perception and cognition. The visual cortex is an ideal system for this enterprise, as retinotopic maps vary significantly in size^37,43^ and their topological organization^10^. However, obtaining retinotopic maps with the precision required for large-scale studies of individual differences is practically impossible, as it requires considerable expertise in stimulus design and specialized data analysis. Despite the existence of multiple large-scale brain imaging datasets of the order of thousands of individuals (including the ABCD dataset^42^, UK Biobank^44^, and HCP Young Adults^41^), currently there exists only a single dataset with retinotopic mapping of the order of hundreds of individuals^26^. This limits our ability to uncover new insights into the functional organization of the visual cortex^10,13^ and its link to perception^11,12,45^. To address this challenge, we leveraged the tight structure-function coupling in the visual cortex. This led to the development of *deepRetinotopy toolbox*, a robust and accessible application for predicting retinotopic maps from brain anatomy at scale, enabling new insights into the functional organization of the human visual cortex.

We found that *deepRetinotopy* accurately predicted polar angle and eccentricity maps across datasets, demonstrating its robustness to differences in imaging sites, magnetic field strength, and scanner. Our method relies solely on curvature maps as input, which are derived from cortical meshes and thus inherently robust to potential intensity variations in the T1w images required for cortical surface reconstruction^46^. In addition, we found that although prediction performance for pRF size maps was high for the HCP test set, performance declined across new datasets. This finding is consistent with known cross-dataset variability in these parameters^47^, by which pRF size is less reliable than polar angle and eccentricity, and pRF size estimates vary considerably depending on the visual stimuli used and fMRI preprocessing pipeline. Nonetheless, polar angle and eccentricity are the primary parameters needed for visual area parcellation and topographic analyses, the key applications of retinotopic mapping.

Having demonstrated the generalizability of our approach, we showed how predicted retinotopic maps can be leveraged to automatically infer visual area boundaries. We found that predicted maps can replace empirical measurements to generate individual-specific early visual area boundaries, providing a scalable alternative to empirical mapping for parcellation in large-scale studies. Moreover, when compared to the gold standard of human manual annotations, we found that automated segmentation can fully replace manual work for V1 segmentation but requires further improvement to reach expert-level segmentation in V2 and V3. Given that we paired *deepRetinotopy toolbox* with an existing Bayesian model of retinotopy to automatically delineate boundaries, substantial improvements are likely to be expected with improvements in the Bayesian modeling approach or by employing new convolutional neural network-based segmentation models^48^.

Finally, we deployed our toolbox to investigate the biological underpinnings of visual cortex organizational differences across age groups, a question that requires sample sizes unattainable with empirical retinotopic mapping. First, we demonstrated that predicted maps capture known asymmetries in visual field sampling in V1^11,13^ (Supplementary Figure 7 and Figure 5), validating that our approach captures genuine organizational principles. Then, we applied *deepRetinotopy* to over 11,000 brain scans (ABCD: n = 9,947; HCP Young Adult: n = 1,113) and found age-related differences in how the primary visual cortex samples the visual field, consistent with prior findings from a small sample study^19^. Importantly, because our predictions derive from cortical geometry alone, these findings reveal that age-related changes in cortical folding may reshape functional organization and exemplify how *deepRetinotopy* enables discoveries at a population scale.

Although our toolbox can generalize and predict retinotopic organization across multiple visual areas from brain anatomy, we focused our performance evaluations on early visual areas (V1-3). Our model’s ability to generate accurate predictions is determined largely by the quality of the training data, which is reflected to some extent in the variance explained from the original pRF analysis^26^. To ensure our models generate better predictions in regions with more reliable pRF estimates (as per the training data), we weighted the loss function by the fraction of variance explained from the pRF solution of each vertex. Note that higher variance explained regions coincide with early visual areas. Despite that, our toolbox still predicted smooth and realistic maps throughout the visual cortex, in over 20 visual areas. However, because the reliability of the training data is lower in higher-order visual areas, at the edge of the stimulus aperture, and in the foveal confluence, we expect that prediction performance may be lower in these regions. Future work could leverage new stimuli tailored for the activation of these regions to improve both the quality of the training data and performance assessment.

While *deepRetinotopy* accurately predicts retinotopic organization in healthy young adult populations, the same cannot be said for other populations with unusual retinotopic organization. For example, in individuals with albinism, V1 exhibits an overlapping representation of the contralateral and ipsilateral hemifields owing to excessive decussation at the optic chiasm^49^, which *deepRetinotopy* would be unable to predict. Moreover, if visual experience changes the spatial tuning of receptive fields across development, *deepRetinotopy* would also be unable to predict experience-dependent organizational changes that are not reflected in cortical anatomy. The reason the group difference in cortical VMA found here has a smaller effect size than the one recently identified^19^ is unclear and could be due to methodological differences or underlying biological explanations. Specifically, the significant difference in cortical VMA identified in the previous study may primarily result from receptive field remapping. A longitudinal retinotopic mapping study could help distinguish between the effects of cortical geometry and receptive field changes on cortical VMA.

Our findings demonstrate how *deepRetinotopy* can provide new insights into the structure-function coupling in the human visual cortex by leveraging large-scale imaging datasets. Our approach offers a standardized framework for both cross-study and cross-individual comparisons, as our methodology is invariant to magnetic field strength, scanner, and acquisition site. This means that data from multiple cohorts can be aggregated and used for normative modeling^50^. Moreover, because our approach can be leveraged at scale, large datasets can be used to enhance statistical power to detect small effects in group comparisons, cross-sectional and longitudinal studies, enabling hypothesis generation and guiding future empirical work.

More broadly, this work establishes a general framework for leveraging structure-function relationships to predict functional organization in other parts of the brain. Future work could spin off our approach to create new predictive models for somatotopic and tonotopic mapping in somatosensory and auditory cortices. Extending our framework to predict topographic organization of more abstract information dimensions, including mental representations of natural objects^51^ as well as semantic representations^52^, would further demonstrate how structure-function coupling principles generalize throughout the brain.

## Methods

### DeepRetinotopy Toolbox

Our toolbox (https://github.com/felenitaribeiro/deepRetinotopy_TheToolbox/) integrates standard neuroimaging software (*FreeSurfer* 7.3.2 and *Connectome Workbench 1.5.0*) for anatomical MRI data preprocessing and a revisited deep-learning model for predicting retinotopic maps^18^ at the individual level. These components are packaged into Docker and Singularity/Apptainer software containers^30^, which can be easily downloaded and are available on Neurodesk^31^. Our toolbox is modular and consists of three main processing steps (Figure 1b):

1. **Input data preprocessing:** Cortical surface meshes are typically derived from preprocessed structural MRI images (T1w) using FreeSurfer (http://surfer.nmr.mgh.harvard.edu/). Starting from FreeSurfer’s output, i.e., the ‘white’ and ‘pial’ surfaces, we generate the midthickness surface using FreeSurfer’s command *mris_expand* or a faster Python-based function that loads both surface meshes as GIFTI files, extracts the vertex coordinates from each surface, and computes the midthickness surface by averaging the corresponding vertex coordinates between the white and pial surfaces. The faster approach has been set as the default one for inference. We then apply FreeSurfer’s *mris_curvature* to the midthickness surface to determine the mean curvature map. These curvature maps are then resampled to the standard HCP ‘32k_fs_LR’ surface space using *Connectome Workbench*.
2. **Retinotopic maps generation:** We leverage a geometric deep learning model^18^ for retinotopic mapping, to predict individual-specific retinotopic maps from curvature maps. To ensure generalizability and ample utility, we revisited previous work^18^ and trained the models to predict retinotopic maps solely from curvature maps generated with *FreeSurfer*. Our current toolbox enables the prediction of polar angle, eccentricity, and pRF size maps from brain anatomy.
3. **Output data processing:** Finally, predicted retinotopic maps are then resampled to the native space of the anatomical data. Specifically, predicted maps are resampled from the standard HCP ‘32k_fs_LR’ surface space to the native space of each individual’s anatomical data using *Connectome Workbench*.

### Datasets

#### Training dataset

All models were trained using a subset of the Human Connectome Project (HCP) 7T Retinotopy dataset, which has been comprehensively described elsewhere^26^. In brief, structural and functional (retinotopic mapping) MRI data were acquired from 181 participants (109 females, age 22-35) following the HCP protocol^26,41^. All participants had normal or corrected-to-normal visual acuity. Structural image acquisition included T1w and T2w structural scans at 0.7 mm isotropic resolution on a customized Siemens 3T Connectom scanner. White and pial cortical surfaces were reconstructed from the structural scans using the HCP pipeline^28^. In preparation for model training, we used FreeSurfer to generate midthickness surfaces for all individuals by expanding the white surface outwards using *mris_expand* and to compute mean curvature maps from the midthickness cortical surfaces with *mris_curvature*. Finally, the mean curvature maps are resampled to the HCP 32k fs_LR standard surface space using the Connectome Workbench command line interface.

Whole-brain retinotopic mapping fMRI data were acquired using a Siemens 7T Magnetom scanner at a resolution of 1.6 mm isotropic and 1 s TR. Retinotopic mapping stimuli were constrained to a circular region with a diameter of 16° and comprised rotating wedges, expanding and contracting rings, and bars of different orientations moving across different directions in the visual field. The retinotopic mapping experiment consisted of 6 runs with different stimulus configurations: one run with contracting rings, one with expanding rings, one run with counterclockwise rotating wedges, one with clockwise rotating wedges, and two identical runs with moving bars. Ring stimuli consisted of 8 cycles of a ring expanding away from the center or contracting towards the center with a period of 32s. Wedges stimuli consisted of 8 cycles of 90° wedges rotating across the visual field counterclockwise or clockwise with a period of 32s. Finally, each bar stimuli run consisted of bars (width=2°) with different orientations (4 orientations) moving across different directions in the visual field. Participants viewed visual stimuli via a back-projection screen using an angled mirror mounted on the head coil. The retinotopic mapping data were processed using the HCP pipeline^28^, which involved correction for gradient distortion, head motion, and EPI image distortion; nonlinear registration of the native volume to MNI space; timeseries mapping from the MNI volume space to the surface space; timeseries resampling from the native surface space to the 32k_fs_LR HCP standard surface space; and denoising for spatially specific structured noise. The data produced (in CIFTI format) by the pipeline consists of 91,282 grayordinates: 32,492 cortical vertices per hemisphere and 26,298 subcortical voxels with approximately 2 mm spatial resolution. All data are publicly available on BALSA (https://balsa.wustl.edu/).

PRF modeling was performed to estimate retinotopic maps from empirical retinotopic mapping fMRI data^26,53^. Essentially, this modeling procedure estimates the spatial sensitivity profile within the visual field to which a grayordinate is responsive (i.e., its receptive field). For this, the fMRI time series elicited by the retinotopic mapping stimuli described above are modeled as a linear function of the portion of the visual stimulus that overlaps with a parameterized model of the pRF at a given location in the visual field^20^. The pRF is typically modeled as a two-dimensional (2D) isotropic Gaussian function with three parameters: *x*₀ and *y*₀, which define the center of the 2D isotropic Gaussian (i.e., the pRF center location), and *σ*, which represents the Gaussian spread, or the population receptive field size. The modeled timeseries is then obtained by computing the dot product between the stimulus aperture time series and a 2D Gaussian (representing the pRF) and then convolving it with a canonical hemodynamic response function. The HCP dataset employed a variation of the traditional pRF model, called the Compressive Spatial Summation model^53^, using analyzePRF, a MATLAB toolbox (http://cvnlab.net/analyzePRF/). For each individual subject, the HCP dataset includes three separate model fits: one fit using all six runs of visual stimuli as described previously (fit 1), a second fit using only the first half of each of the six runs (fit 2), and the third fit using only the second half of each of the six runs (fit 3).

Prior to model training, the 181 participants from the HCP 7T retinotopy dataset were separated into three datasets as in our previous work^18^: training (161 participants), development (10 participants), and test (10 participants). Each subset was used for training the neural network, model selection, and benchmarking, respectively.

#### Generalizability datasets

To assess model generalizability, we collated a comprehensive set of datasets acquired under different conditions, namely: imaging sites, imaging scanners, and visual stimuli used for retinotopic mapping experiments. These datasets include: the New York University (**NYU**) Retinotopy dataset^33,54^, the **Stanford** child and adult checkboard retinotopy dataset^19,55,56^, the Center for Human Neuroscience (**CHN**) retinotopic mapping dataset^57,58^, and **New Zealand** dataset^59^ (available on request). Table 2 provides the main features of each of these datasets. Data acquisition, preprocessing, and pRF modeling from each dataset were comprehensively described elsewhere, and all derivative data (preprocessed data as well as pRF model fits) are openly available via OpenNeuro data repositories, except for the New Zealand dataset. Each one of these datasets obtained retinotopic maps using different algorithmic implementation of the pRF model^20^. The NYU and the Stanford datasets both employed *vistasoft* (Vista Lab, Stanford University; https://github.com/vistalab/vistasoft) for pRF model fits, which is a MATLAB toolbox. The CHN dataset employed custom MATLAB software. The New Zealand dataset employed SamSrf (https://github.com/samsrf/samsrf), a MATLAB toolbox. We did not perform any additional processing of the structural data apart from the ones performed via our toolbox (with default settings). All retinotopic maps were resampled to the 32k_fs_LR HCP standard space and polar angle maps were transformed to ensure consistency of angle representation for generalizability assessment.

**Table 2.**
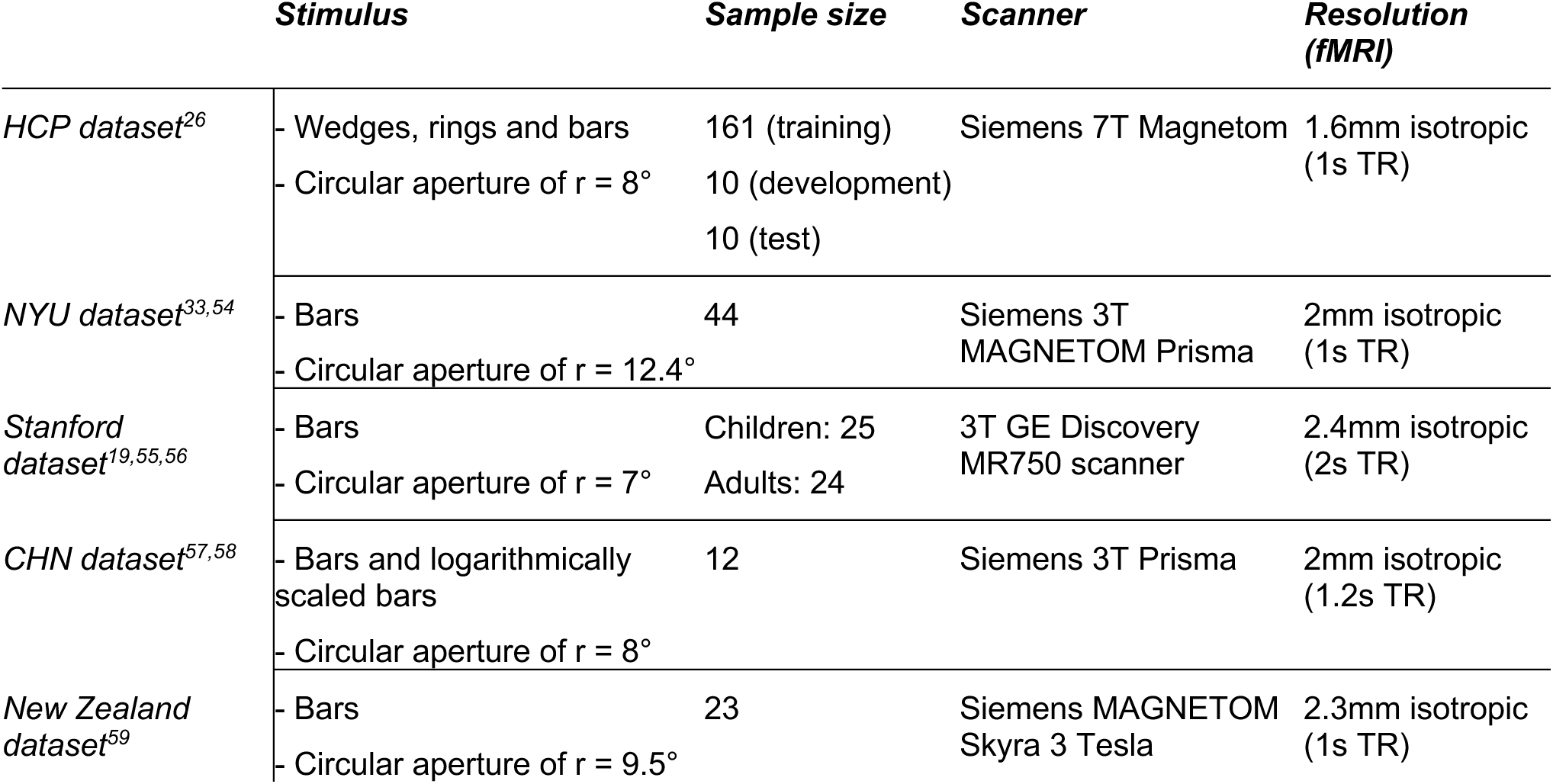
Summary of relevant information from datasets used in this work.

### Regions of interest

Models were trained to predict retinotopic organization within a region of interest (ROI) encompassing all visual areas from the Wang et al. surface-based probabilistic atlas^8^, except the frontal eye field (FEF) due to its spatial discontinuity with the other visual areas. This region of interest contained 3,267 vertices in the left hemisphere and 3,219 vertices in the right hemisphere. Throughout the manuscript, we focus our analyses on early visual areas, which include vertices from V1, V2, and V3 and their foveal confluence, selected from the intersection of the full visual cortex ROI and early visual area masks.

### Model training and selection

Comprehensive description of training data, model architecture, and training has been provided elsewhere^18^. Briefly, training data (n=161 participants) included: the mean curvature maps (input data) resampled to the HCP 32k fs_LR standard surface space; a template data frame on which the curvature maps are represented, i.e., a template cortical surface (S1200_7T_Retinotopy181.L(R).midthickness_MSMAll.32k_fs_LR.surf.gii); expected output data, which was either polar angle, eccentricity, or pRF size. To avoid issues with the discontinuity in polar angle maps due to its cyclical nature (i.e., 360°= 0°), we shifted the polar angle values from the left hemisphere so that the point of wrap-around (from 360° to 0°) was positioned at the horizontal meridian in the contralateral hemifield.

Our geometric deep learning model consists of a spline-based convolutional neural network^60^ (SplineCNN) developed to perform convolution operations on surfaces using PyTorch Geometric^61^. We did not change our previously reported model architecture, except for the number of input features in the first layer. As in our previous work, our models’ learning objective was to reduce the difference between predicted retinotopic map and ground truth (i.e., the empirically derived map). This mapping objective is measured by the smooth L1 loss function, where the difference between empirically derived and predicted parameter was weighted by the individual-specific explained variance (R^2^) from the pRF modeling procedure^26^. Models were implemented using Python 3.8.13, PyTorch 2.4.1, PyTorch Geometric 2.6.1, and CUDA 12.1. Training was performed on a high-performance computing cluster using an NVIDIA H100 GPU.

We trained 5 distinct models per retinotopic map and hemisphere using different random initializations, totalling 30 models. Owing to the large model weights’ files (∼465MB each) and slow inference speed using CPU, we made our toolbox available with a single instance model per retinotopic map and hemisphere, totalling 6 models, which minimizes the software container size and inference speed. Note, however, that all pre-trained models’ weights are publicly available on an OSF repository (https://osf.io/ermbz/). The final models were selected based on the error and individual variability metrics, such that higher interindividual variability across predicted retinotopic maps would not come at the cost of higher prediction error. The error was estimated as the difference between the predicted and the empirically derived angle values in a vertex-wise manner and averaged across all participants from the HCP development dataset (n=10). Individual variability was determined by the difference between a specific predicted map and each other predicted map in the development dataset in a vertex-wise manner and averaged across all combinations (9 combinations) and then averaged across participants. To estimate the difference between two angles, we used the smallest difference between two angles as our metric, given by:

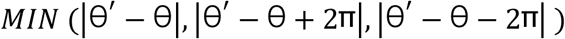

Supplementary Figure 8 shows the error and individual variability across model seeds, retinotopic maps, and hemisphere, for the individuals within the HCP development set.

### Benchmarking

To benchmark our method, we compared our approach with two other methods for predicting retinotopic organization from brain anatomy. These are: a model with the same architecture as ours using both curvature and myelin maps as input features^18^ and the anatomical template of retinotopy^15^, which we refer to as **deepRetinotopy-beta** and **Benson2014**, respectively. DeepRetinotopy-beta consists of the same geometric deep learning model architecture as our approach, but takes both curvature and myelin maps as input features. Thus, we trained one model per retinotopic map and hemisphere using the curvature maps derived from the midthickness surface, as previously described, and myelin maps provided by the HCP dataset, which were determined by T1w/T2w images ratio. To train these models we used the same training data as in our current approach. Benson2014 consists of an anatomical template of retinotopy to which individual participant structural data is registered, yielding retinotopic predictions at the individual-level^15^. We obtained predictions with the latter approach using *Neuropythy*^9^ (https://github.com/noahbenson/neuropythy).

Performance across methods was determined by computing both the correlation score and error between the empirically estimated and the predicted retinotopic maps. Correlation score was determined as either the Pearson correlation (for eccentricity and pRF size maps) or the circular correlation (polar angle maps) for each individual in the HCP test set (n = 10). These measures were obtained using Python packages *Scipy*^62^ and *Astropy*^63^. The error was determined as either the smallest angular difference for polar angle maps (as in the previous section) or the absolute difference for eccentricity and pRF size maps, calculated in a vertex-wise manner and averaged across vertices within a given region of interest. All analyses were restricted to vertices with variance explained > 10% and within V1-3 to focus on reliably measured retinotopic responses.

To determine the upper and lower boundaries of models’ performance, we estimated the noise ceiling and error floor. The noise ceiling is the best possible performance achievable by any model and can be estimated using split halves of the data. Specifically, we estimated the noise ceiling via the Spearman-Brown formula, given as:

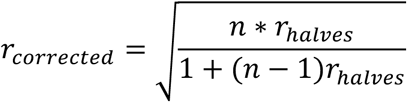

where *r*_*halves*_ is the correlation between retinotopic maps estimated using only the first half (fit 2 as provided by the HCP dataset) and only the second half (fit 3) of the empirical retinotopic mapping data, and *n* represents the number of splits (n = 2) being combined to estimate the noise ceiling. Finally, the error floor was determined as the difference between the retinotopic maps derived from each split half, calculated as either the smallest angular difference (for polar angle maps) or the absolute difference (for eccentricity and pRF size maps).

### Generalizability

We assessed model generalizability both qualitatively and quantitatively using four distinct datasets, as described above. Qualitative assessment was performed via visual inspection of hexbin plots, which represents the density of data points in a two-dimensional space. These plots were generated by first vectorizing empirically derived and predicted maps and filtering out vertices outside of V1-3 and below a variance explained threshold of 10% of the empirically derived pRF parameters. Next, vectorized maps were concatenated across participants. These data were then represented as hexbin plots, where empirically derived parameters were shown along the y-axis and the predicted ones along the x-axis.

Quantitatively, we determined the correlation score between the empirically derived and predicted maps, using either the Pearson correlation (for eccentricity and pRF size maps) or the circular correlation (polar angle maps), for each individual and averaged across individuals within each dataset. To assess statistical significance, we used a spatial autocorrelation-preserving null model (‘spin test’)^35^. Null distributions were derived from *n* = 1000 ∗ *sample size* maps derived from random rotations to spherical projections of empirically derived maps from the HCP test set using *BrainSpace*^64^. These null maps were then correlated with their corresponding predicted maps to determine a null correlation distribution for statistical testing.

### Visual area segmentation

Visual area labels were estimated using the Bayesian model of retinotopy^9^, which combines anatomical priors with empirical observations to generate individual-specific retinotopic maps and visual area boundaries. The model uses a retinotopic template (prior) and retinotopic maps derived from some minutes of fMRI data (observation) to produce individual-level predictions. We use either the empirically estimated or the predicted retinotopic maps as observations to derive visual area labels automatically. Moreover, the Bayesian model weights observations based on variance explained, a proxy for data quality: higher variance explained of the empirical data by the pRF model leads to greater influence of the observation relative to the prior. Given that variance explained can only be estimated when empirical retinotopic mapping data are available, we generate proxy weight maps when *deepRetinotopy*’s predicted parameters are used as observations for the Bayesian model inference.

First, we compared how similar the automatically generated visual area labels were to manually drawn labels. To do so, we used the test and development sets from the HCP combined, as manually drawn labels by four anatomists^37^ are openly available on the Open Science Framework^65^ (n = 16 with all manual annotations available). Here, we used the mean variance explained across all training participants as the weight map to be used with the predicted retinotopic maps. We then computed the degree of overlap between manually drawn and automatically generated early visual area labels, which was determined by the Dice score. Second, we estimated the impact of the weighting on the automatic estimation of the visual area labels by using an alternative weight map with ones everywhere (maximum weight to the observation; see Supplementary Figure 9). We similarly computed the degree of overlap between manually drawn and automatically generated early visual area labels. To establish the theoretical ceiling for automated segmentation performance, we computed inter-rater agreement by calculating Dice scores between all pairs of manual annotations, averaged across anatomists and participants.

### Estimating V1 surface area dedicated to sample different portions of the visual field

To estimate the cortical surface area of V1 representing different portions of the visual field, we established an automated framework inspired by an earlier work^19^. First, we combine V1 and V2 masks, derived from the Wang et al. surface-based probabilistic atlas^8^, with their foveal confluence to ensure V1 was fully contained within our region of interest. We then selected vertices within a specific eccentricity ranges (0-6°). The polar angle maps were then used to mask vertices representing different portions of the visual field:

- the upper vertical meridian (UVM), which includes vertices representing 90°± ***d***;

- the lower vertical meridian (LVM), which includes vertices representing 270°± ***d***;

- the right horizontal meridian (RHM), with vertices representing angles between 0°+ ***d*** and 360°- ***d***;

- the left horizontal meridian (LHM), which includes vertices representing 180°± ***d***, where ***d*** was equal 45°.

To minimize the inclusion of vertices falling within each of these ranges but that are in unexpected regions of the visual cortex, we also use Wang et al.’s probabilistic atlas ROIs to restrict upper/lower vertical meridians to ventral/dorsal portions of our ROI, respectively. Finally, we estimate the cortical surface area of each of these wedge-ROIs with custom Python code, using the cross-product method for triangle area calculation from the midthickness surface in *fsnative* space. The estimated cortical surface areas are then used to calculate the cortical vertical-meridian asymmetry (VMA) and the cortical horizontal-vertical anisotropy (HVA). In both cases, the upper and lower vertical surface areas were divided by a factor of 2 to account for the additional vertices from V2 included in our masking procedure. Cortical VMA is given by:

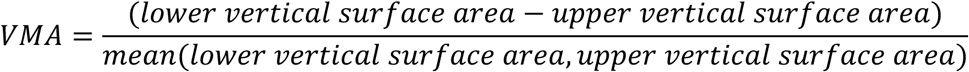

where a positive cortical VMA index indicates a bigger surface area dedicated to representing the lower vertical portion of the visual field than the upper portion.

Similarly, the cortical HVA is given by:

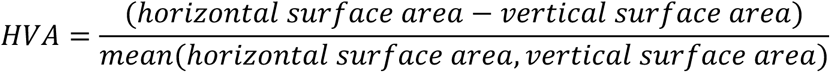

where a positive cortical HVA index indicates a bigger surface area dedicated to representing the horizontal portion of the visual field than the vertical portion.

### Visualization

Surface plots were generated using Nilearn^66^, a Python module for fast statistical learning on neuroimaging data. Graphs were generated using a combination of Seaborn^67^, Pandas^68^, and Matplotlib^69^ functionalities.

## Code availability

The *deepRetinotopy toolbox* source code is available on GitHub (https://github.com/felenitaribeiro/deepRetinotopy_TheToolbox) as well as the code required to reproduce all our experiments (https://github.com/felenitaribeiro/deepRetinotopy_validation). The toolbox is available as Docker and Singularity containers and via Neurodesk^31^.

## Acknowledgement

FLR acknowledges support through the European Union’s Horizon Europe research and innovation funding program under the Marie Skłodowska-Curie Actions project ID 101146996. FLR and SB acknowledge funding by an Australian Research Council Linkage grant (LP200301393) awarded to SB. RS acknowledges support through a doctoral scholarship from the German Academic Scholarship Foundation as well as the European Union’s Horizon Europe research and innovation program, grant 101039712. NCB acknowledges support from the National Eye Institute, grant 1R01EY033628. DL acknowledges funding by the Austrian Science Fund (FWF) [doi.org/10.55776/P35583]. MNH was supported by the ERC Starting Grant COREDIM (ERC-2021-STG-101039712), a LOEWE Start Professorship by the Hessian Ministry of Higher Education, Research, Science and the Arts, and the Deutsche Forschungsgemeinschaft (German Research Foundation, DFG) under Germany’s Excellence Strategy (EXC 3066/1 “The Adaptive Mind”, Project No. 533717223). Open access funding provided by Max Planck Society.

The training data were provided by the Human Connectome Project, WU-Minn Consortium (Principal Investigators: David Van Essen and Kamil Ugurbil; 1U54MH091657) funded by the 16 NIH Institutes and Centers that support the NIH Blueprint for Neuroscience Research; and by the McDonnell Center for Systems Neuroscience at Washington University. This work was supported by resources provided by The University of Queensland Research Computing Centre’s Bunya supercomputer^70^. We thank Torin Bambridge-Lozan for supporting with an early prototype of the inference pipeline through the UQ AI Collaboratory Summer Internship Program. We thank the Neurodesk team for providing support with tool packaging, and Tomas Knapen and Sander van Bree for helpful discussions. The authors acknowledge that artificial intelligence tools (including ChatGPT, Claude and Grammarly) were used for brainstorming, text editing, and proofreading.

## Author contribution

**Fernanda L. Ribeiro:** Conceptualization, Methodology, Software, Validation, Formal analysis, Investigation, Data curation, Writing – Original Draft, Review & Editing, Visualization, Supervision, Project administration, Funding acquisition. **Robert Satzger:** Software, Formal Analysis, Investigation, Writing – Review & Editing. **Felix Hoffstaedter:** Methodology, Software, Formal Analysis, Data Curation, Writing - Review & Editing. **Christian Bürger:** Software, Investigation, Writing - Review & Editing. **Peer Herholz:** Software, Writing - Review & Editing. **David Linhardt:** Data curation, Writing – Review & Editing. **Noah C. Benson:** Software, Writing - Review & Editing. **D. Samuel Schwarzkopf:** Software, Data curation, Writing – Review & Editing. **Alexander M. Puckett:** Conceptualization, Writing - Review & Editing. **Steffen Bollmann:** Software, Resources, Writing – Review & Editing, Supervision, Funding acquisition. **Martin N. Hebart:** Conceptualization, Methodology, Resources, Writing – Review & Editing, Supervision, Funding acquisition.

## Supplementary information

**Supplementary Figure 1.**
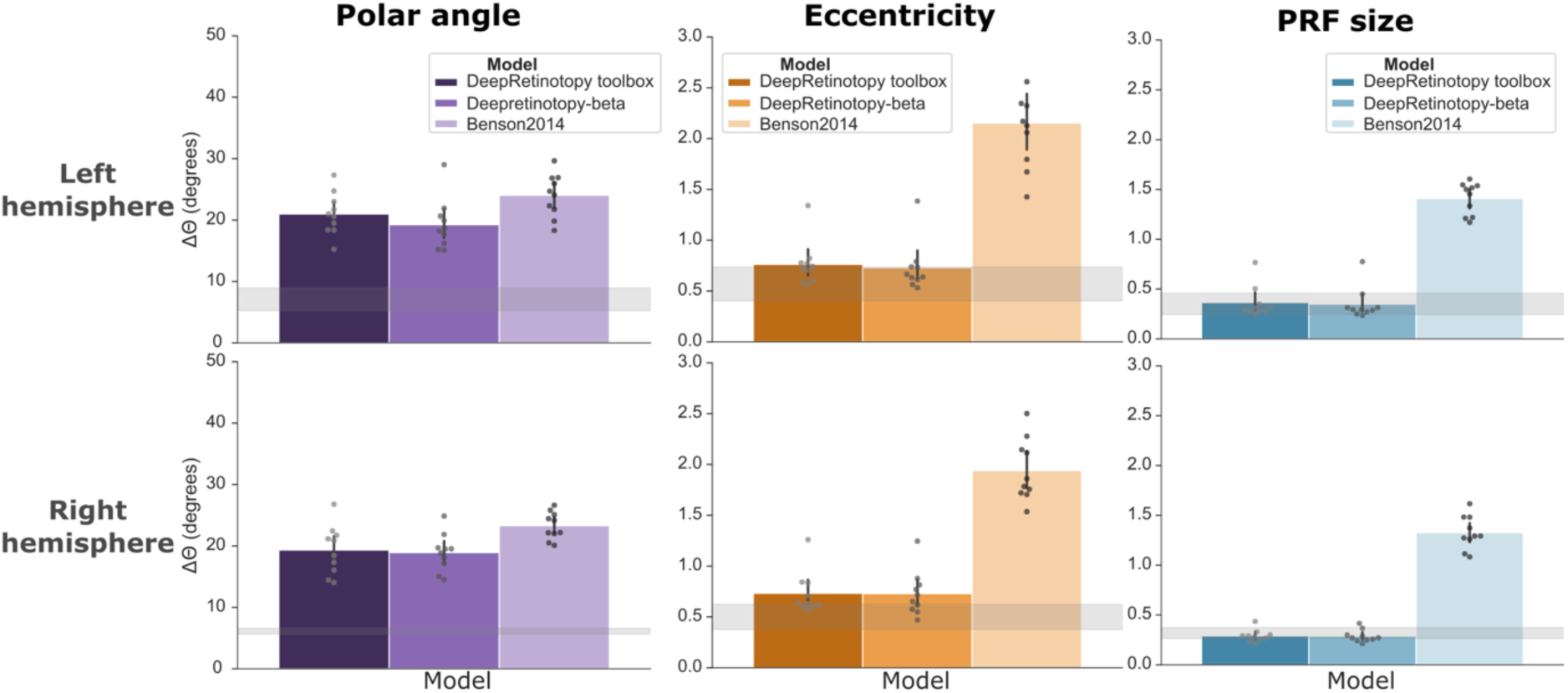
Model benchmarking with an error measure. Bar plots represent th average error between predicted and empirically derived maps across all participants in the HCP test set. Retinotopic maps were vectorized and only vertices within early visual areas (V1-3) and above a 10% variance explained threshold were used to estimate the error given as the smallest angular difference for polar angle maps or the absolute difference for eccentricity and pRF size maps. The gray shaded area represents the 95% confidence interval of the error floor, i.e., the difference between the retinotopic maps derived from each split half.

**Supplementary Figure 2.**
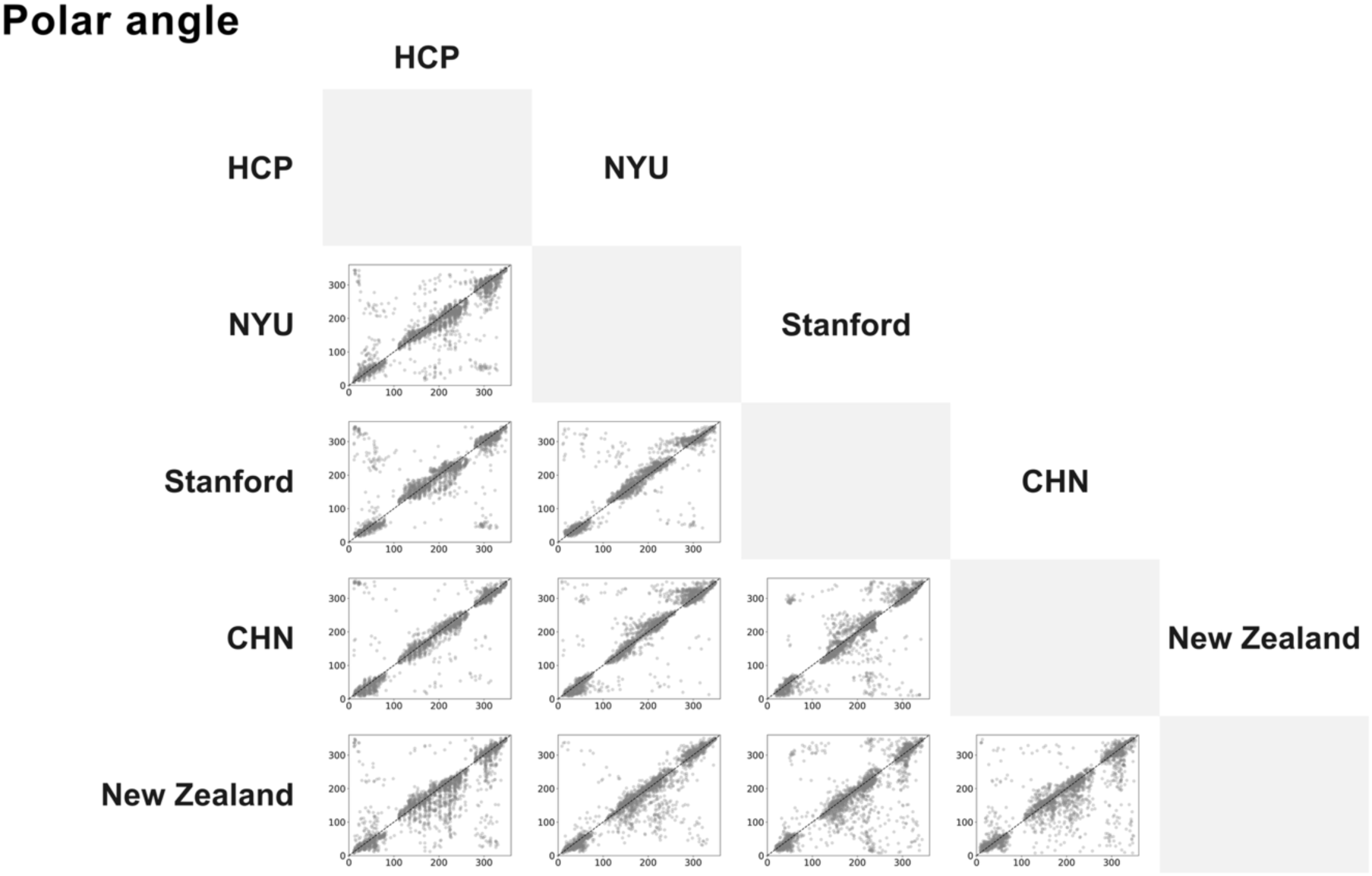
Cross-dataset comparison of polar angle maps of the visual cortex. Scatter plots comparing vertex-wise median parameters across participants from each test dataset as in Figure 2. Each data point represents a vertex in the fs_LR_32k surface space within V1-3. Data was aggregated across hemispheres, and we also applied a variance explained threshold of 15% based on datasets along the Y axis.

**Supplementary Figure 3.**
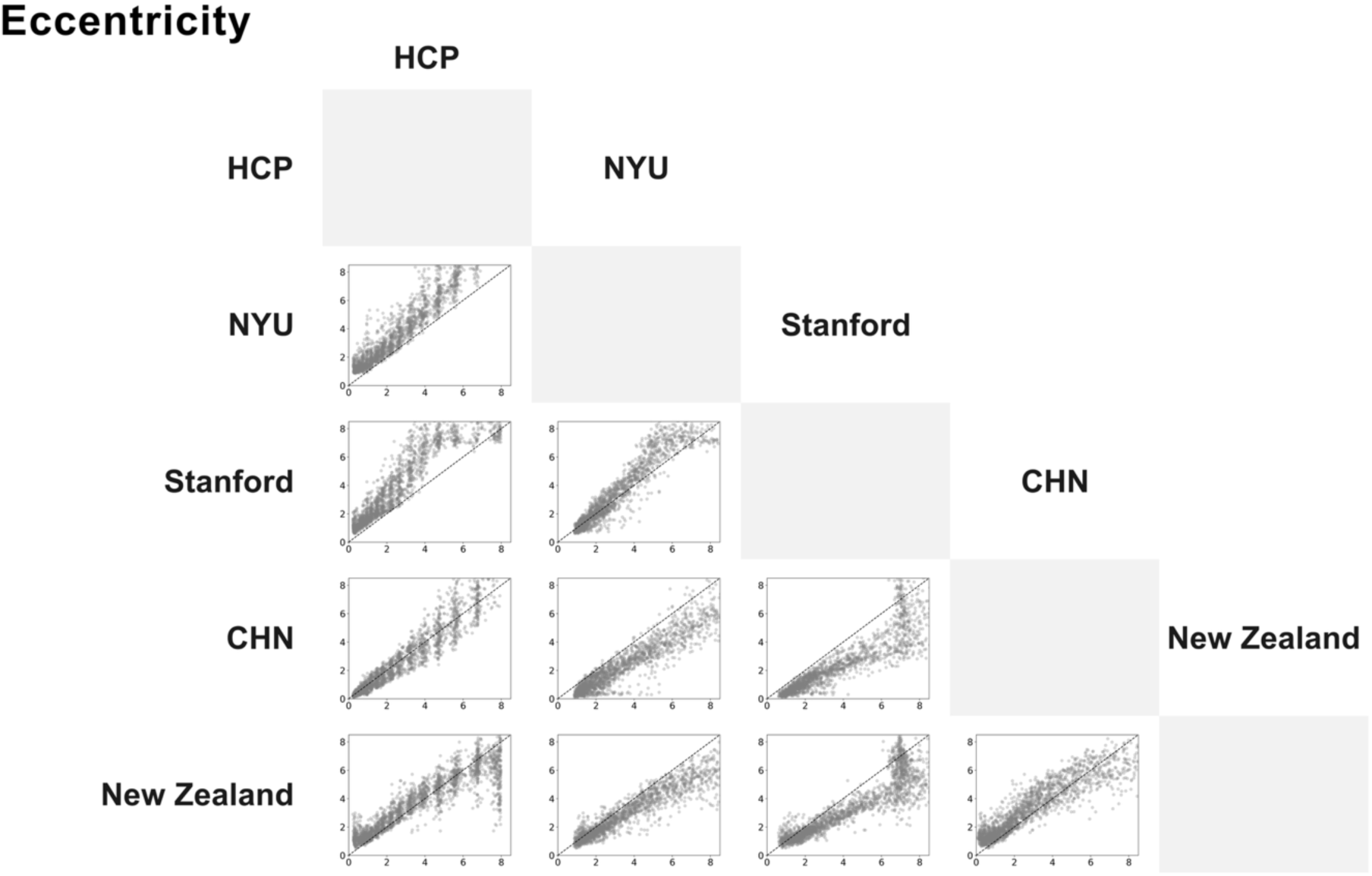
Cross-dataset comparison of eccentricity maps of the visual cortex. Scatter plots comparing vertex-wise median parameters across participants from each test dataset as in Figure 2. Each data point represents a vertex in the fs_LR_32k surface space within V1-3. Data was aggregated across hemispheres, and we also applied a variance explained threshold of 15% based on datasets along the Y axis.

**Supplementary Figure 4.**
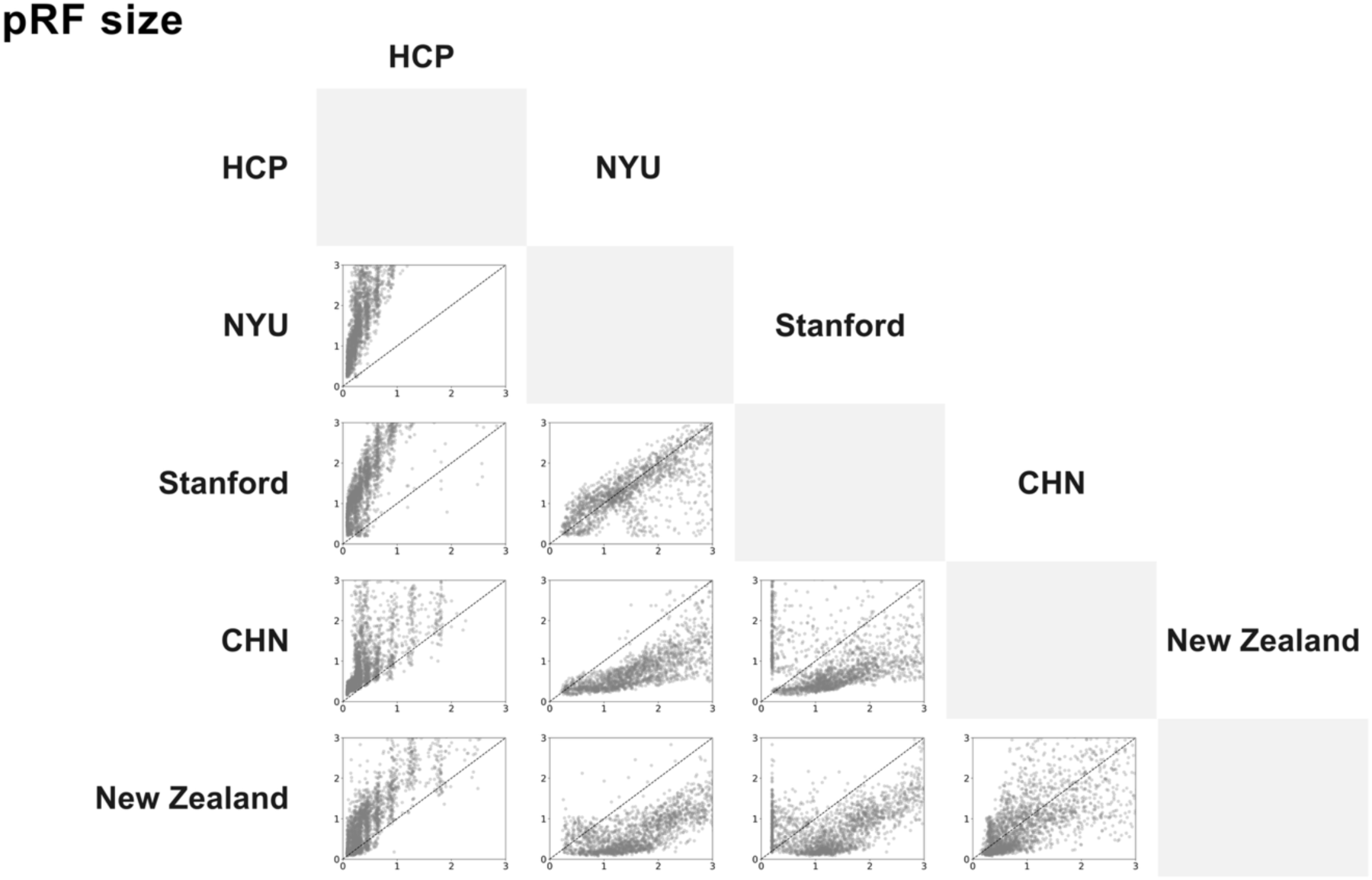
Cross-dataset comparison of pRF size maps of the visual cortex. Scatter plots comparing vertex-wise median parameters across participants from each test dataset a in Figure 2. Each data point represents a vertex in the fs_LR_32k surface space within V1-3. Data was aggregated across hemispheres, and we also applied a variance explained threshold of 15% based on datasets along the Y axis.

**Supplementary Figure 5.**
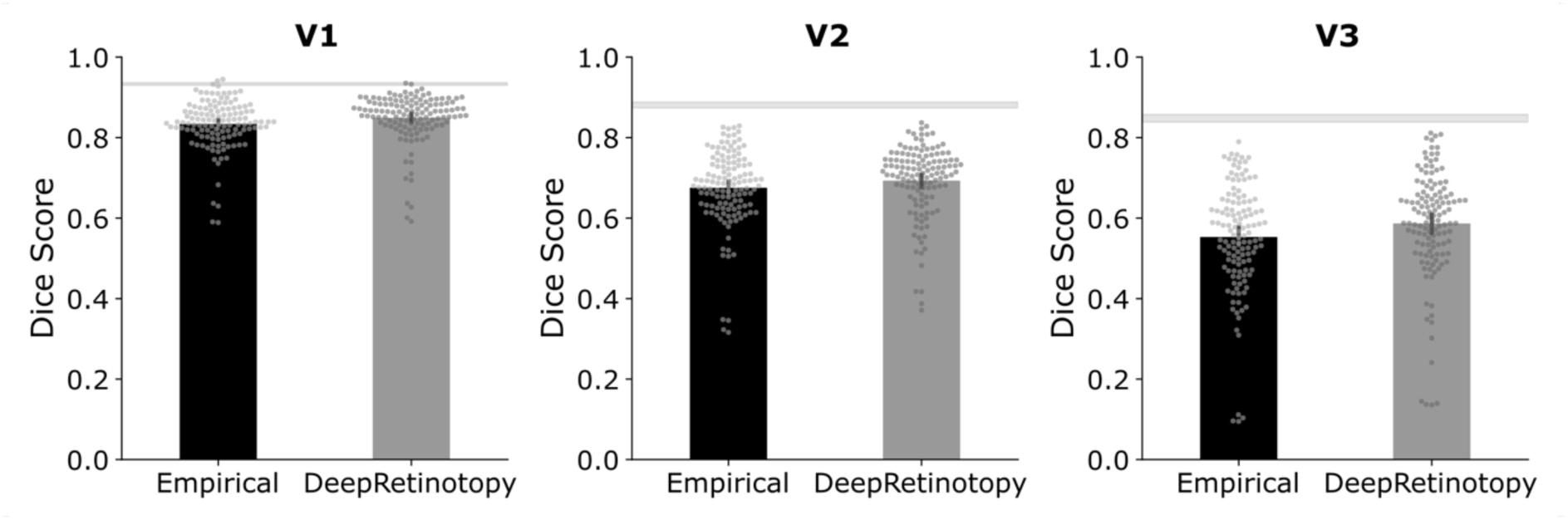
Automated visual area segmentation performance (non-normalized). Performance is shown across visual areas. Performance was estimated as the degree of overlap between manually drawn and automatically generated early visual area labels, for which data from both hemispheres were combined. Error bars correspond to the 95% confidence interval. The gray shaded area represents the noise ceiling, i.e., the 95% confidence interval of the DICE scores between all pairs of manual annotations, across anatomists and participants.

**Supplementary Figure 6.**
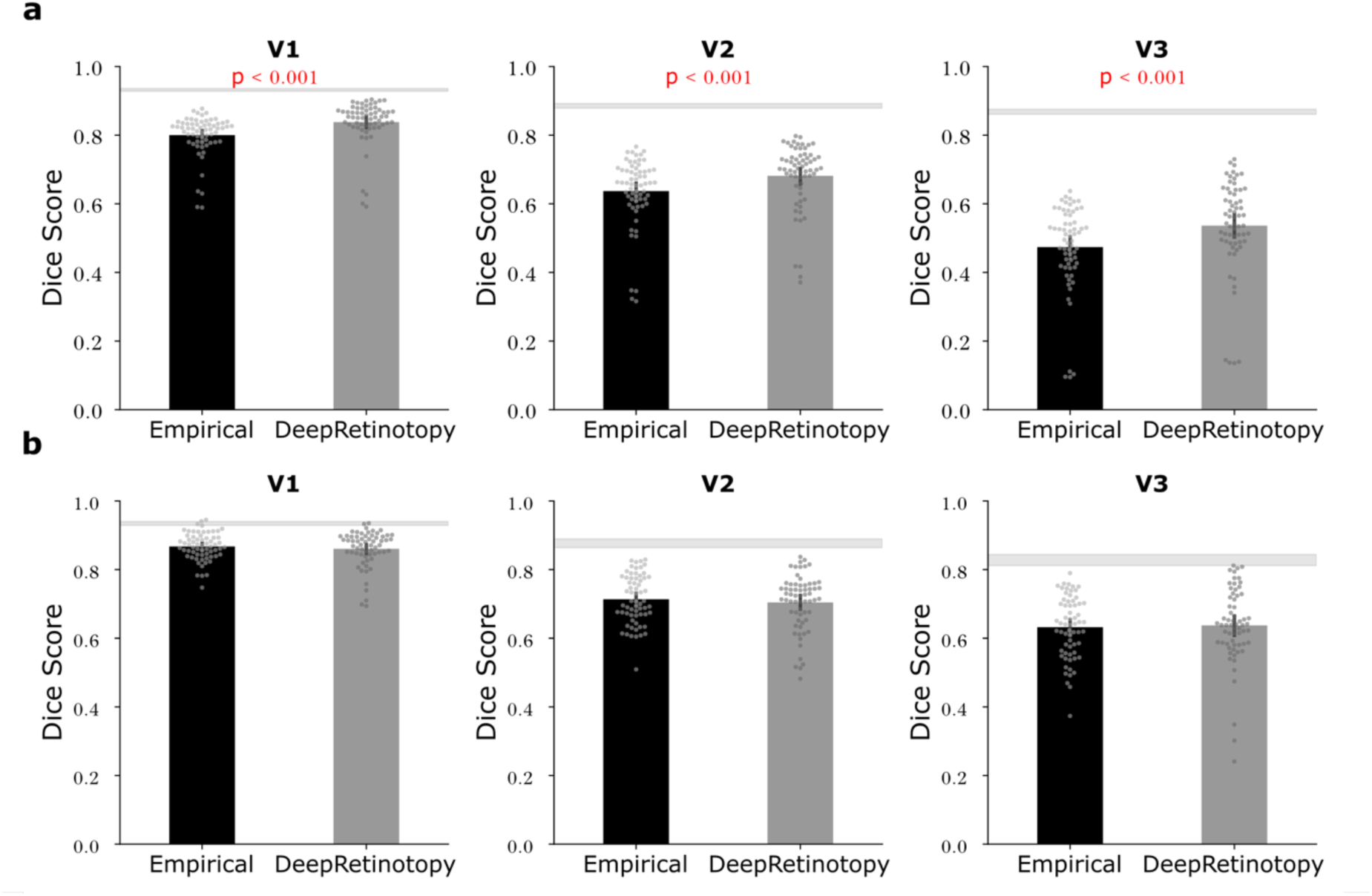
Segmentation performance across visual areas per hemisphere. Plots show the Dice score across visual areas for right (**a**) and left (**b**) hemispheres separately. The gray shaded area represents the noise ceiling, i.e., the 95% confidence interval of the Dice scores between all pairs of manual annotations, across anatomists and participants.

**Supplementary Figure 7.**
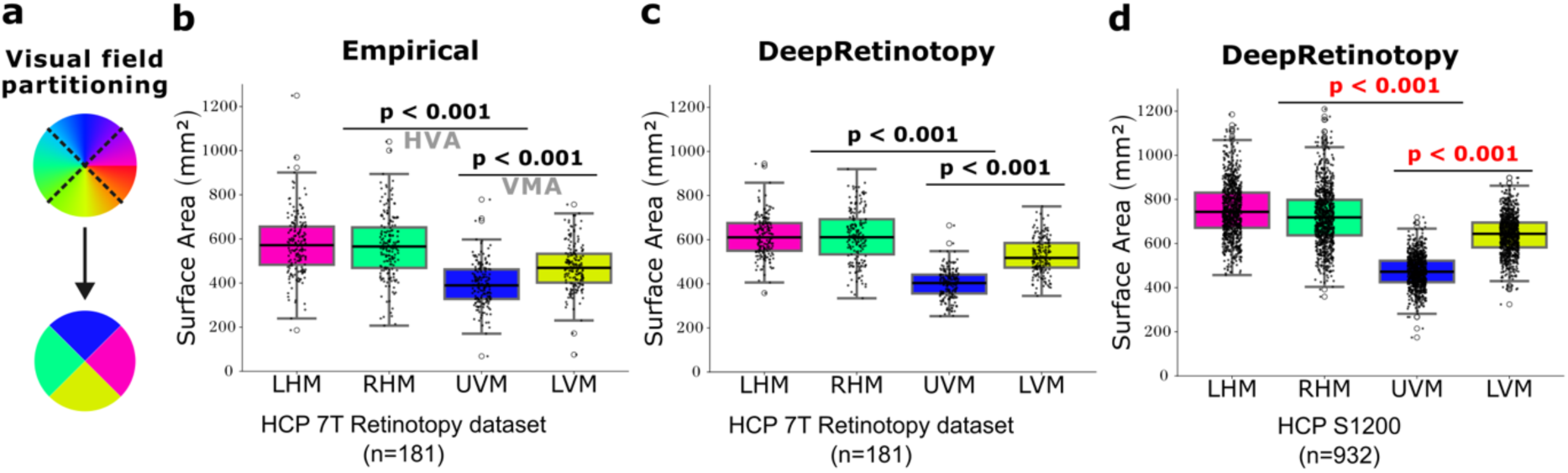
Polar angle asymmetries for V1 surface area. **a,** Diagram shows visual field partitioning used for determining wedge-ROIs to estimate the cortical surface area dedicated to representing the left (pink) and right(green) horizontal meridians, and upper (blue) and lower (yellow) vertical meridians. Group-level V1 surface area measures from wedge-ROIs are shown for empirically derived (**b**) and predicted retinotopic maps (**c**) using the HCP 7T Retinotopy dataset (n=181), and for predicted retinotopic maps using the HCP young adult dataset (**d**; n = 932, excluding the individuals with retinotopic mapping data). Black data points indicate individual measurements. The top and bottom bounds of each box represent the 75th and 25th percentiles, respectively. LHM: left horizontal meridian; RHM: right horizontal meridian; UVM: upper vertical meridian; LVM: lower vertical meridian; HVA: horizontal-vertical anisotropy; VMA: vertical-meridian asymmetry.

**Supplementary Figure 8.**
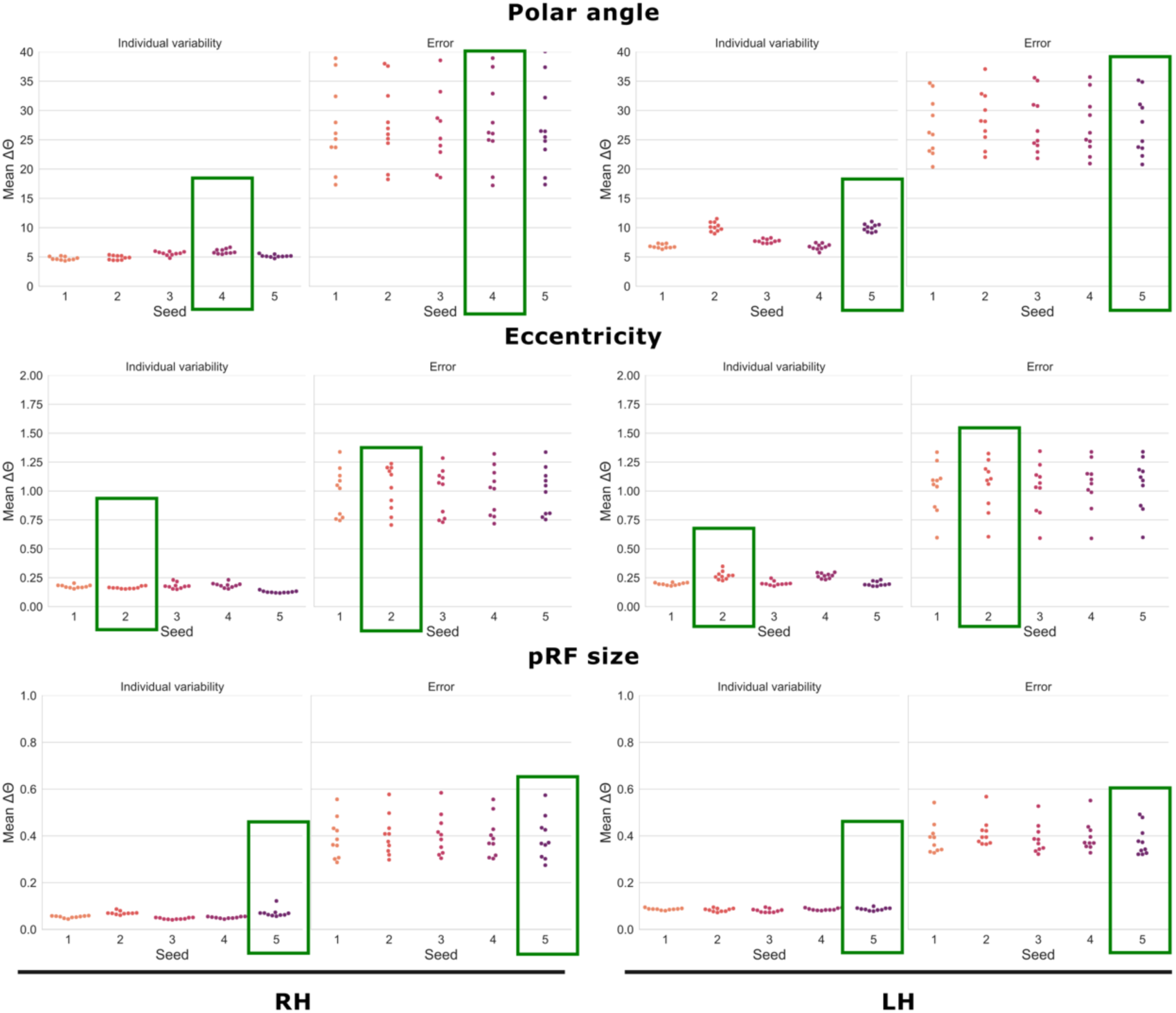
Model performance in the development set (n = 10) across different random initializations. We trained 5 distinct models per retinotopic map (polar angle, eccentricity, and pRF size) and hemisphere (left: LH; right: RH) using different random initializations, totaling 30 models. Owing to the large model weights’ files (∼465MB each) and slow inference speed using CPU, we made our toolbox available with a single instance model per retinotopic map and hemisphere, totaling 6 models, which minimizes the software container size and inference speed. The green rectangles highlight the selected models. Note we based our selection on both the individual variability and error scores.

**Supplementary Figure 9.**
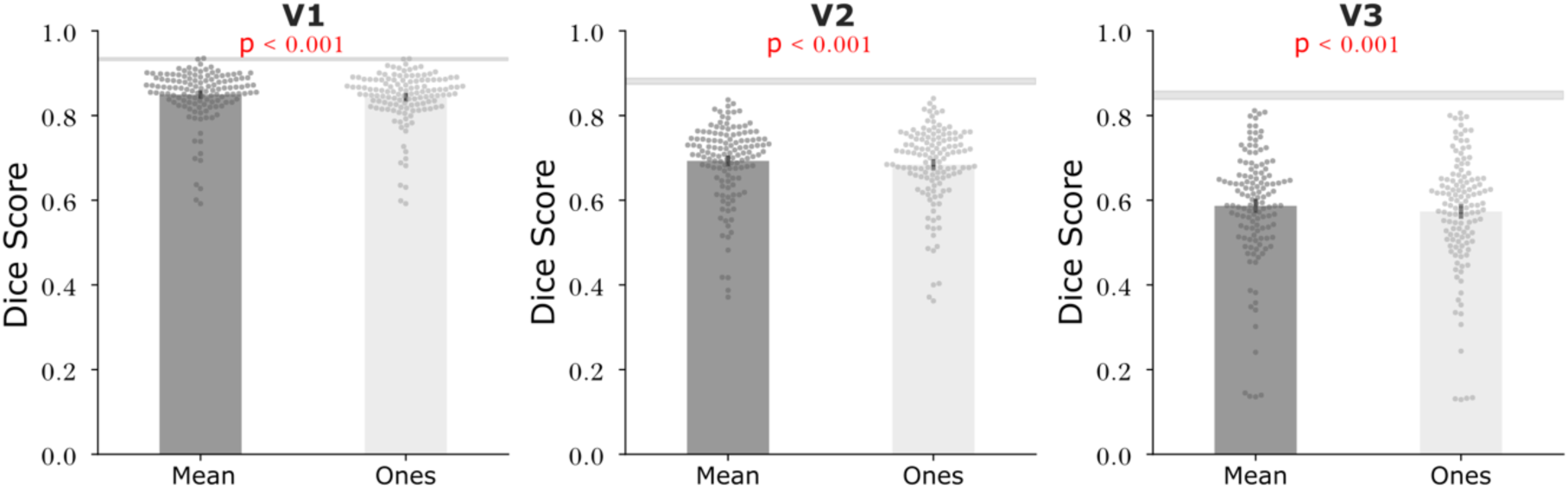
Impact of weighting on automated visual area segmentation performance. Segmentation performance is shown across visual areas, with data from both hemispheres combined. We compared two weighting approaches for the Bayesian model: the mean variance explained across all *deepRetinotopy* training participants *versus* uniform maximum weighting (ones everywhere, giving maximum weight to the predicted observations). The variance explained- based weighting achieved higher segmentation performance (V1: mean Dice score = 0.85; V2: 0.69; V3: 0.59) compared to uniform weighting (V1: 0.84; V2: 0.68; V3: 0.57). Statistical significance was assessed using two-tailed paired t-tests. The gray shaded area represents the noise ceiling, .e., the 95% confidence interval of the Dice scores between all pairs of manual annotations, averaged across anatomists and participants.

**Supplementary Table 1.** Mean correlation scores between the empirically derived and predicted maps across datasets and hemispheres. Correlation scores were determined as the Pearson correlation for eccentricity and pRF size maps and the circular correlation for polar angle maps. LH: left hemisphere; RH: right hemisphere.

| *Dataset* | *Hemisphere* | *Retinotopic map* | *Correlation* | *p-value* |
| --- | --- | --- | --- | --- |
| *HCP* | LH | Polar angle | 0.8335 | 0.00e+00 |
|  |  | Eccentricity | 0.8825 | 0.00e+00 |
|  |  | PRF size | 0.6807 | 0.00e+00 |
| *NYU* |  | Polar angle | 0.6907 | 0.00e+00 |
|  |  | Eccentricity | 0.7593 | 0.00e+00 |
|  |  | PRF size | 0.6868 | 0.00e+00 |
| *Stanford* |  | Polar angle | 0.6736 | 0.00e+00 |
|  |  | Eccentricity | 0.6898 | 0.00e+00 |
|  |  | PRF size | 0.6124 | 0.00e+00 |
| *CHN* |  | Polar angle | 0.8103 | 0.00e+00 |
|  |  | Eccentricity | 0.8638 | 0.00e+00 |
|  |  | PRF size | 0.4855 | 0.00e+00 |
| *New Zealand* |  | Polar angle | 0.637 | 0.00e+00 |
|  |  | Eccentricity | 0.7275 | 0.00e+00 |
|  |  | PRF size | 0.533 | 0.00e+00 |
| *HCP* | RH | Polar angle | 0.8415 | 0.00e+00 |
|  |  | Eccentricity | 0.9009 | 0.00e+00 |
|  |  | PRF size | 0.7614 | 0.00e+00 |
| *NYU* |  | Polar angle | 0.6803 | 0.00e+00 |
|  |  | Eccentricity | 0.7444 | 0.00e+00 |
|  |  | PRF size | 0.7038 | 0.00e+00 |
| *Stanford* |  | Polar angle | 0.6492 | 0.00e+00 |
|  |  | Eccentricity | 0.7015 | 0.00e+00 |
|  |  | PRF size | 0.6223 | 0.00e+00 |
| *CHN* |  | Polar angle | 0.8304 | 0.00e+00 |
|  |  | Eccentricity | 0.859 | 0.00e+00 |
|  |  | PRF size | 0.4038 | 0.00e+00 |
| *New Zealand* |  | Polar angle | 0.5509 | 0.00e+00 |
|  |  | Eccentricity | 0.6966 | 0.00e+00 |
|  |  | PRF size | 0.5038 | 0.00e+00 |

